# Unlocking Predictive Power: A Machine Learning Tool Derived from In-Depth Analysis to Forecast the Impact of Missense Variants in Human Filamin C

**DOI:** 10.1101/2023.08.05.552086

**Authors:** Michael Nagy, Georg Mlynek, Julius Kostan, Luke Smith, Dominic Pühringer, Philippe Charron, Torsten Bloch Rasmussen, Zofia Bilinska, Mohammed Majid Akhtar, Petros Syrris, Luis R Lopes, Perry M Elliott, Mathias Gautel, Oliviero Carugo, Kristina Djinović-Carugo

## Abstract

Cardiomyopathies, diseases of the heart muscle, are a leading cause of heart failure. An increasing proportion of cardiomyopathies have been associated with specific genetic changes, such as mutations in *FLNC*, the gene that codes for filamin C. Altogether, more than 300 variants of *FLNC* have been identified in patients, including a number of single point mutations. However, the role of a significant number of these mutations remains unknown. Here, we conducted a comprehensive analysis, starting from clinical data that led to identification of new pathogenic and non-pathogenic *FLNC* variants. We selected some of these variants for further characterization that included studies of *in vivo* effects on the morphology of neonatal cardiomyocytes to establish links to phenotype, and the *in vitro* thermal stability and structure determination to understand biophysical factors impacting function. We used these findings to compile vast datasets of pathogenic and non-pathogenic variant structures and developed a machine-learning-based neural network (AMIVA-F) to predict the impact of single point mutations. AMIVA-F outperformed most commonly used predictors both in disease related as well as neutral variants, approaching ∼80% accuracy. Taken together, our study documents additional *FLNC* variants, their biophysical and structural properties, and their link to the disease phenotype. Furthermore, we developed a state-of-the-art web-based server AMIVA-F that can be used for accurate predictions regarding the effect of single point mutations in human filamin C, with broad implications for basic and clinical research.

## Introduction

Heart failure remains a major global health concern that affects millions of people worldwide, leading the Global Burden of Diseases study to define it as a global epidemic in 2017 (1). Among others, cardiomyopathies, diseases of the heart muscle that affect mechanical and/or electrical function of the heart, represent one of the leading causes of heart failure (2). Approximately 20% to 40% of nonischemic cardiomyopathies are caused by detectable genetic changes identified in over 70 different genes associated with heart muscle development and function (3). One of the genes that is frequently found mutated in patients with familial cardiomyopathies is *FLNC*, a gene that codes for filamin C (FlnC) (4). For example, mutations in *FLNC* have recently been associated with hypertrophic cardiomyopathies (HCM), restrictive cardiomyopathies (RCM), arrhythmogenic cardiomyopathy (ACM) and dilated cardiomyopathy (DCM) (4–9). Thus far, more than 300 *FLNC* variants have been described in the literature (4), although not every variant is pathogenic. Therefore, the increasing number of *FLNC* variants potentially associated with cardio-and other types of myopathies highlights the need for developing bioinformatic tools for assessing their pathogenicity.

To be accurate and effective, these bioinformatic tools need to integrate the accumulated knowledge about the *in vivo* function and the *in vitro* properties of FlnC. Functionally, FlnC is a member of filamin family of proteins that serve to connect components of the actin cytoskeleton to sarcolemma, and the only filamin found in cardiac muscles. FlnC localizes to sarcolemma, myotendinous junctions, intercalated discs and Z-discs, and functions at the boundaries between adjacent sarcomeres, which are the basic contractile unit of the muscle (10). To achieve this function, FlnC interacts with a plethora of other proteins, either Z-disc components (e.g. myotilin, calsarcins, myopodin, LDB3/ZASP, Xin), signalling molecules or sarcolemma associated proteins (e.g. integrin ß1, sarcoglycan delta (reviewed in (11)). Furthermore, FlnC is recruited to sarcomeric lesions together with aciculin/phosphoglucomutase-5 to accomplish its role in myofibril stabilization and early myofibril repair processes (12–14). Structurally, filamins, including FlnC, are composed of an N-terminal actin-binding domain (ABD), followed by 24 immunoglobulin (Ig)-like domains (Ig1-Ig24) (10,15). Filamin Ig-like domains are organized *via* intricate inter-domain interactions (16–20) that are the basis of the force-/mechano-sensing and ligand binding, with dynamic interactions between Ig domains of the C-terminal region, in the playing a mechanosensing function (**Figure 1B**) (21,22). In addition, contrary to non-muscle filamins (FlnA and FlnB), FlnC contains a unique feature, an 82-amino acid insertion in Ig20 (insertion region, IR), with a recently proposed role in modifying the fine molecular details of mechanosensing or/and acting as an interaction hub for Z-disc proteins (11). Many reported *FLNC* variants are single point mutations distributed across all Ig-like domains; thus, making predictions regarding how they affect structure and function of FlnC, as well as predicting the extent of their pathogenicity, has been challenging.

**Figure 1.**
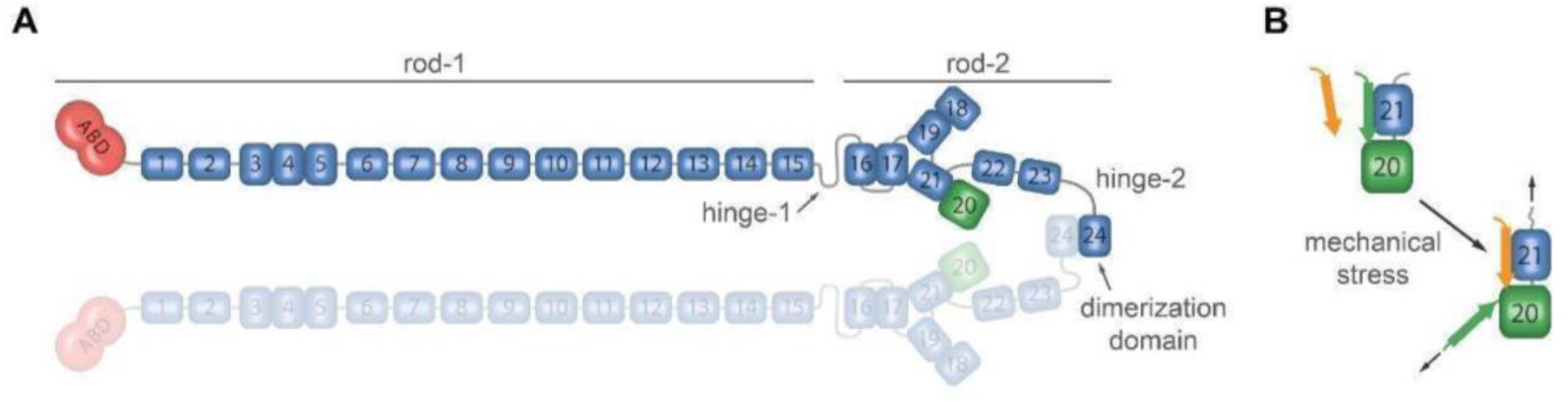
Domain structure and mechanosensing mechanism of filamin. (**A**) Domain structure of FlnC dimer. ABD is depicted in red, Ig-domains in blue, domain 20 hosting the insertion region of 82 aa in green. Inter-and intra-molecular domain pairing is schematically presented. Hinge-1 is spliced out during myogenesis (10,15). (**B**) Proposed mechanism of mechanosensing and ligand binding regulation (16,22,23). Under unstrained conditions β-strand A of Ig20 covers the ligand-binding site present in domain Ig21. Mechanical force will lead to domain pair opening, exposing therefore the cryptic binding site in Ig21 to interactions with binding partners (16,22,23).

Here, we present a comprehensive study that encompasses clinical data, which led to identification of new *FLNC* variants, and cell-based studies, biophysical and structural analysis of both known and novel FlnC mutants. These studies extended our understanding of the functional and structural effect of FlnC mutations. We combined the new insights with the information available in the literature, to design a machine learning algorithm AMIVA-F that can be used as a bioinformatics tool for predicting the pathogenicity of FlnC missense variants. We confirmed that AMIVA-F outperforms currently available tools. Furthermore, analysis of available information on pathogenic mutants combined with structural data allowed us to draw conclusions about the relationship between amino acid conservation amongst human FlnC Ig domains and pathogenicity.

## Results

### Patient clinical data, demographics and family histories

We report data on eight patients, six of whom carry a pathogenic *FLNC* variant, and two who carry a neutral variant. Overall, the cohort included both men (2 of 8) and women (6 of 8), all except one of European origin (one woman in this cohort is of Afro-Caribbean origin). In this cohort, all patients who bore pathogenic variants exhibited features of various cardiomyopathies (see below for details). No clinical skeletal myopathy was noticed in these patients. Below we describe each patient, their demographic information, and provide summaries of associated clinical data and family histories. The patient information is organized by the location of their point mutation, from N to C-terminus of FlnC.

<u>Patient #1</u> is a female proband, who received a diagnosis of dilated cardiomyopathy (DCM) at 23 years of age. She had an implantable cardioverter-defibrillator (ICD) at 23 (appropriate shocks for ventricular fibrillation), left ventricular ejection fraction (EF) 22% at presentation and had heart transplantation (Htx) four years later. Patient #1 family history includes an affected brother who died suddenly with arrhythmogenic right ventricular cardiomyopathy (ARVC) described in the postmortem, a maternal side first cousin with dilated cardiomyopathy (DCM), known to carry the same variant and with an ICD in situ, and affected mother and maternal aunt, both dying suddenly at 36 and 43 years of age, respectively. In both cases, the results of the post-mortem described left-dominant ARVC. We showed that Patient #1 is the carrier of variant c.245T.G that results in p.Met82Lys point mutation. This point mutation is located within the ABD of FlnC. Currently, this mutation is classified as “uncertain significance” by the American college of Medical Genetics and Genomics (ACMG).

<u>Patient #2</u> is a white male proband, with a diagnosis of restrictive cardiomyopathy (RCM) established at 40 years of age with moderate left ventricular (LV) systolic dysfunction, and presence of atrial tachyarrhythmias. He is in permanent atrial fibrillation (AF). His echocardiogram showed mild concentric left ventricular hypertrophy (LVH), left ventricular ejection function (LVEF) of 40-45% and a markedly enlarged left atrium. Cardiac magnetic resonance (CMR) showed concentric remodelling with a LV maximal wall thickness (MWT) of 13 mm with widespread fibrosis or late gadolinium enhancement (LGE) both in the mid-wall in the septum and sub-epicardially in the basal anterior and mid lateral walls. He has had a primary prevention subcutaneous ICD implanted and a decompensated heart failure (HF) admission during follow-up. He has two children with a diagnosis of RCM and known to carry the same variant, and both have primary prevention ICDs. We determined that Patient #2 carries *FLNC* variant c.4871C.T, resulting in p.Ser1624Leu point mutation. This point mutation is located within immunoglobulin domain 14 (Ig14). Currently, this mutation is classified as “conflicting interpretation of pathogenicity” by ACMG.

<u>Patient #3</u> is a white female, with a diagnosis of hypertrophic cardiomyopathy (HCM). Her brother had a diagnosis of HCM/RCM, with HTx, her mother had HCM and died after HTx, and her mother’s cousin had HCM and also died after HTx. There is no family history of sudden cardiac death (SCD). She was first evaluated when admitted at 43 years of age with symptoms of dyspnoea, presyncope and palpitations. Initial echocardiogram revealed asymmetric septal hypertrophy with a MWT of 17 mm and a dilated left atrium. CMR demonstrated a LVEF of 61% and the presence of limited late gadolinium enhancement in the inferior wall and mid-septum. She remained symptomatic in New York Heart Association (NYHA) class II dyspnoea and in persistent AF over follow-up and had ablation for this at 54 years of age. There was no evidence of nonsustained ventricular tachycardia (NSVT) or sustained VT throughout follow-up, and no evidence of peripheral myopathy, including normal creatine kinase. We discovered that Patient #3 carried the same variant as Patient #2.

<u>Patient #4</u> is a white male proband, with a diagnosis of RCM. He presented at 18 years-old with symptoms of NYHA class II dyspnoea, chest pain, and palpitations. His initial echocardiogram demonstrated normal wall thickness, mildly dilated left atrium, good systolic function, LVEF 75%. Doppler was consistent with restrictive physiology. CMR showed preserved biventricular function and no evidence of LGE. He had one HF admission at 24 years of age and recurrent readmissions due to AF and developed significant pulmonary hypertension that responded favourably to oral magnesium treatment of 1g/day. Since 2012, he has remained ambulatory in NYHA class III/IV heart failure. NSVT was identified on Holter at last evaluation and repeated CMR demonstrated significant left atrial dilatation, heterogeneous midwall LGE, in basal and middle septal segments and anterior wall. He had no peripheral myopathy, including normal levels of serum creatine kinase. He was considered for HTX referral. He is a carrier of *FLNC* variant c.5026G>A, yielding p.Gly1676Arg mutation within Ig15. This variant was identified in ten relatives, all of them with an RCM phenotype of different degrees, with atrial arrhythmias (four in AF). Of these, three cousins died, two of severe heart failure at the age of 61 years and 63 years and another cousin of stroke at age of 56 years. There was no sudden death in the family. Currently, this mutation is classified as “uncertain significance” by the ACMG.

<u>Patient #5</u> is a white female proband, with an initial diagnosis of HCM that later progressed to RCM. She was first seen aged 27, with NYHA class II and a previous transitory ischaemic attack. She had persistent AF at baseline. She was admitted due to HF for the first time aged 38 and her functional class evolved to class IV during follow-up. She had a HTx aged 50. Patient #5 was discovered to carry *FLNC* variant c.6053G.C, resulting in p.Arg2018Pro point mutation located within Ig18. Twelve relatives are known to carry the same variant, nine of them clinically affected, including two who had HTx, one who died of HF and one ICD carrier for secondary prevention. Currently, this mutation is not classified by ACMG.

<u>Patient #6</u> is a white female, with a high ventricular ectopic burden and family history of arrhythmogenic right ventricular cardiomyopathy. Her son died aged 26 with ARVC described on PM. Her paternal grandfather had SCD aged 54. Her paternal second cousin was diagnosed with ARVC and had an ICD implanted after a presentation with VT. Her baseline Holter identified 456 ventricular ectopics (VEs)/24 hours. She had a normal echocardiogram and almost normal CMR in her late 60s, with only minor basal lateral fibrosis on CMR. Most recent Holter monitoring has demonstrated increased ventricular ectopy (819 VEs/24 hour, 9% burden). She does not fulfil criteria for an overt cardiomyopathy phenotype at 72. Patient #6 carries *FLNC* variant c.6173A>G, that causes p.Gln2058Arg mutation in Ig19. Currently, this mutation is classified as “uncertain significance” by the ACMG.

<u>Patient #7</u> is a white female with a family history of SCD during sleep in her son, who died aged 36 and was diagnosed with ARVC post-mortem, including a description of LV subtle fibrofatty infiltration. Her echocardiogram showed mild LVH with MWT of 12 mm and grade 1 diastolic dysfunction but was otherwise normal. Her CMR demonstrated normal biventricular size and function, but with a subtle streak of sub-epicardial LGE in the basal to mid inferior wall. SAECG was negative for late potentials and Holter monitoring did not detect any arrhythmia. Patient #7 is a carrier of *FLNC* variant c.6779A>G, resulting in p.Lys2260Arg point mutation within Ig20. This patient has a subtle cardiomyopathy phenotype aged 79. Currently, this mutation is classified as “conflicting interpretation of pathogenicity” by ACMG.

<u>Patient #8</u> is an Afro-Caribbean female, diagnosed with RCM. She had a family history of RCM in her father, who died aged 55 and, in her sister, who had a HTx aged 22 years and subsequently died. She also had two other sisters with RCM. There was no FH of SCD. She was first evaluated at 36 and was symptomatic with NYHA class III dyspnoea and decompensated HF. She was noted to be in atrial tachycardia and had AF ablation at 38. She had multiple HF admissions and no NSVT or sustained VT. Patient #8 was shown to carry *FLNC* variant c.6892C>T, resulting in p.Pro2298Leu point mutation within Ig20. Currently, this mutation is not classified by ACMG.

Collectively, the analysis of clinical data allowed us to identify 8 new *FLNC* variants associated with disease phenotypes of varying severity. Structurally, the resulting point mutations were distributed across the entire protein, thus providing a good representation of *FLNC* variants previously reported in the literature.

### Impact of mutations p.Met82Arg, p.Ser1624Leu and p.Gly1676Arg in cellular context

To investigate the effects of selected variants on myofibril morphology in a cellular context, we transfected GFP-FlnC bearing specific mutations into neonatal cardiomyocytes and analysed them by fluorescence confocal microscopy. We chose to focus on p.Met82Lys, p.Ser1624Leu and p.Gly1676Arg as these three mutations cover several human disease phenotypes, and map to different domains of FlnC, the ABD, and two Ig-domains (Ig14, Ig15), respectively. These three constructs showed distinctive cellular phenotypes when compared to wild-type FlnC (**Figure 2**). Wild-type FlnC localized predominantly to the Z-disc of transfected cells, where it colocalized with Z-disc titin (**Figure 2**). Furthermore, we observed that actin cytoskeletal structures also contain filamin and titin in a non-striated pattern at the cell periphery and that some cells showed a faint overlay of the FlnC construct around the sarcomere centre, at or above the M-band.

**Figure 2.**
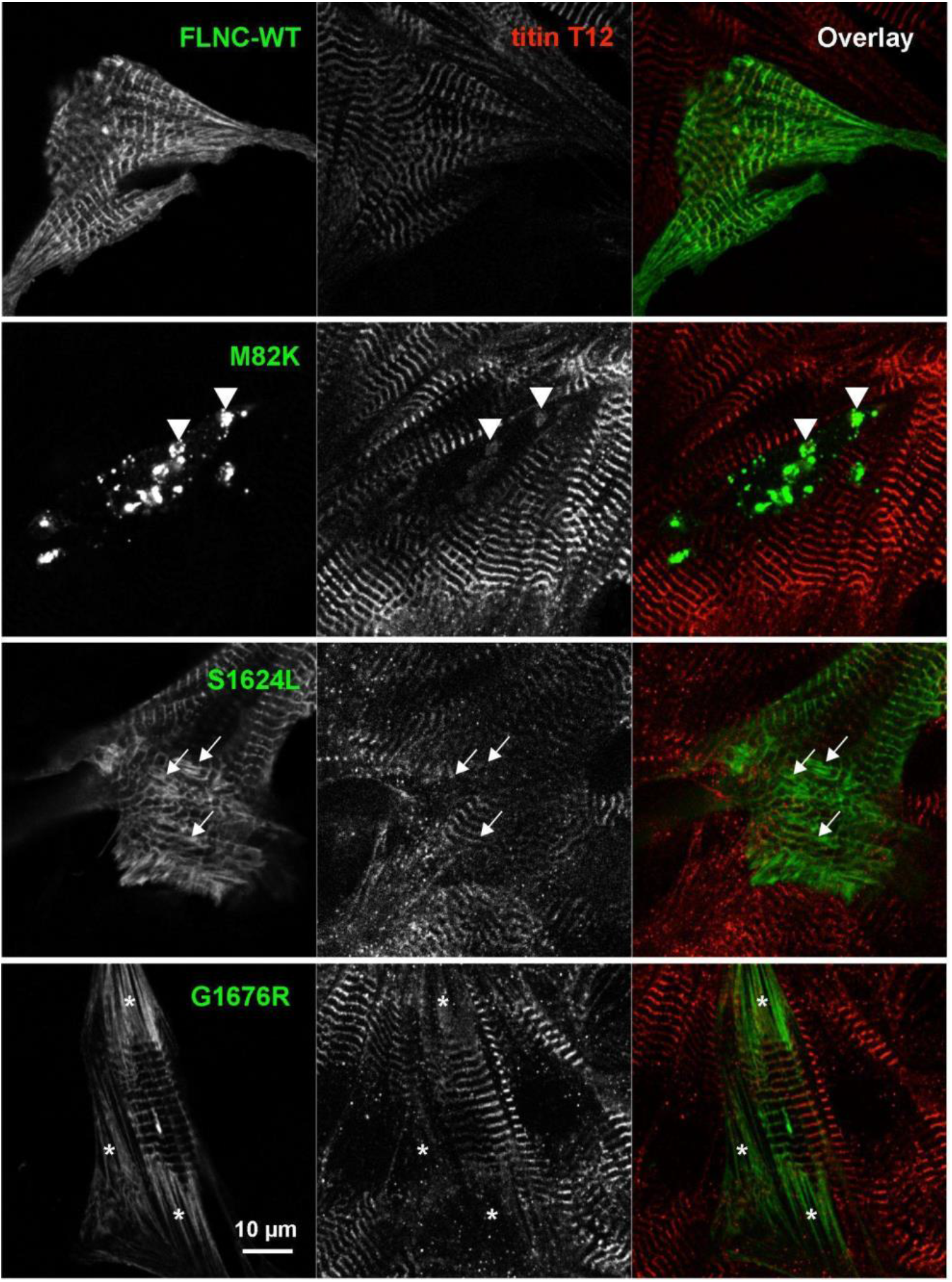
In cell validation of selected variants reported in this study. Neonatal rat cardiomyocytes were transfected with GFP-tagged FlnC constructs to visualise integration into sarcomeres or other structures. Top row: wild-type filamin-C (FlnC-WT) localises to Z-discs where it overlaps with the titin-Z-disc epitope T12. Towards the cell periphery, non-striated myofibrils, presumed precursors of myofibrils, contain both titin and FlnC in an irregular localisation. Second row: the ABD mutant p.Met82Lys (labeled M82K) forms pronounced aggregates (arrowheads) and in some cases leads to the disappearance of sarcomeres, as evidenced by co-localising with Z-disc titin in aggregates. Third row: the mutation p.Ser1624Leu (labeled S1624L) in Ig14 leads to sarcomere disarray and the pronounced formation of Z-disc streaming (arrows). Bottom row: the mutation p.Gly1676Arg (labeled G1676R) in Ig15 leads to reduction of the cross-striated cardiomyocyte area with extensive non-striated areas of cytoskeleton containing titin and FlnC in a stress-fibre like structures (asterisks). Titin T12 red, GFP-FlnC green. Scale bar: 10 µm.

In contrast, cells transfected with the mutant variants displayed a range of abnormalities. The p.Met82Lys variant showed cytoplasmic aggregates in many cells leading to the disappearance of striated titin and its localization to the FlnC-containing aggregates (arrowheads in **Figure 2**), suggesting a dominant effect on sarcomere structure. On the other hand, cardiomyocytes transfected with the FlnC variant p.Ser1624Leu showed a subtler phenotype without strong disruption of sarcomeres. The variant localised to Z-discs but also to long “Z-disc streams” of aberrantly organised sarcomeric structures (arrows in **Figure 2**). Lastly, the cellular effect of the p.Gly1676Arg variant was again distinct. Cardiomyocytes expressing this variant showed a marked reduction of the striated (sarcomeric) actin cytoskeleton, with extensive areas of non-striated actin cytoskeleton dominating the periphery of the cells (asterisks in **Figure 2**). Quantification of the images indicated that the ratio of striated versus non-striated actin cytoskeleton decreased from 0.8 for wild-type transfected cells to 0.48 for the p.Gly1676Arg variant. This observation could indicate a dominant effect on the maturation or maintenance of sarcomeric structures by this variant. Taken together, the cell-based analysis illustrates that each of the disease-associated *FLNC* variants affects appearance of cardiomyocytes in different ways, causing a wide range of effects and perturbations when compared to the wild-type cells.

### Impact of pathogenic variants on structural integrity of FlnC

In order to assess whether the pathogenicity is rooted in the altered stability of the domains, we performed thermal stability and structural analyses. We selected eight variants, four of which are reported in the literature (p.Ile2160Phe (Ig20), p.Trp2164Cys (Ig20), p.Pro2298Ser (Ig20), p.Ser1624Leu (Ig14)), and five of which have been described in the section **Patient clinical data, demographics and family histories** (p.Met82Lys (ABD), p.Ser1624Leu (Ig14), p.Gly1676Arg, (Ig15), p.Gln2058Arg (Ig19) and p.Lys2260Arg (Ig20)) (**<u>Table S1</u>**). All the constructs yielded sufficient amounts of purified protein for biophysical stability characterization, except the ABD p.Met82Lys mutant. Met82Lys ABD did not express in *E. coli*, suggesting that this point mutation causes significant destabilization of the protein fold. This effect is supported by analysis of the ABD structure that shows that Met82 side-chain is embedded within a hydrophobic environment. Thus, Met-to-Lys substitution is expected to cause severe steric clashes and repulsion, leading to destabilization and potential unfolding of the domain (**Figure 3A**). Impaired folding of the ABD may negatively impact its solubility and interactions with actin filaments (F-actin), and thus lead to misslocalization of the protein, which is in agreement with our cell-based observations (**Figure 2**). As mentioned, we were able to express other Ig14-Ig15 and Ig19-Ig21 constructs bearing single point mutations and used differential scanning calorimetry (DSC) to analyze their thermal stability (**<u>Table S1</u>**).

**Figure 3.**
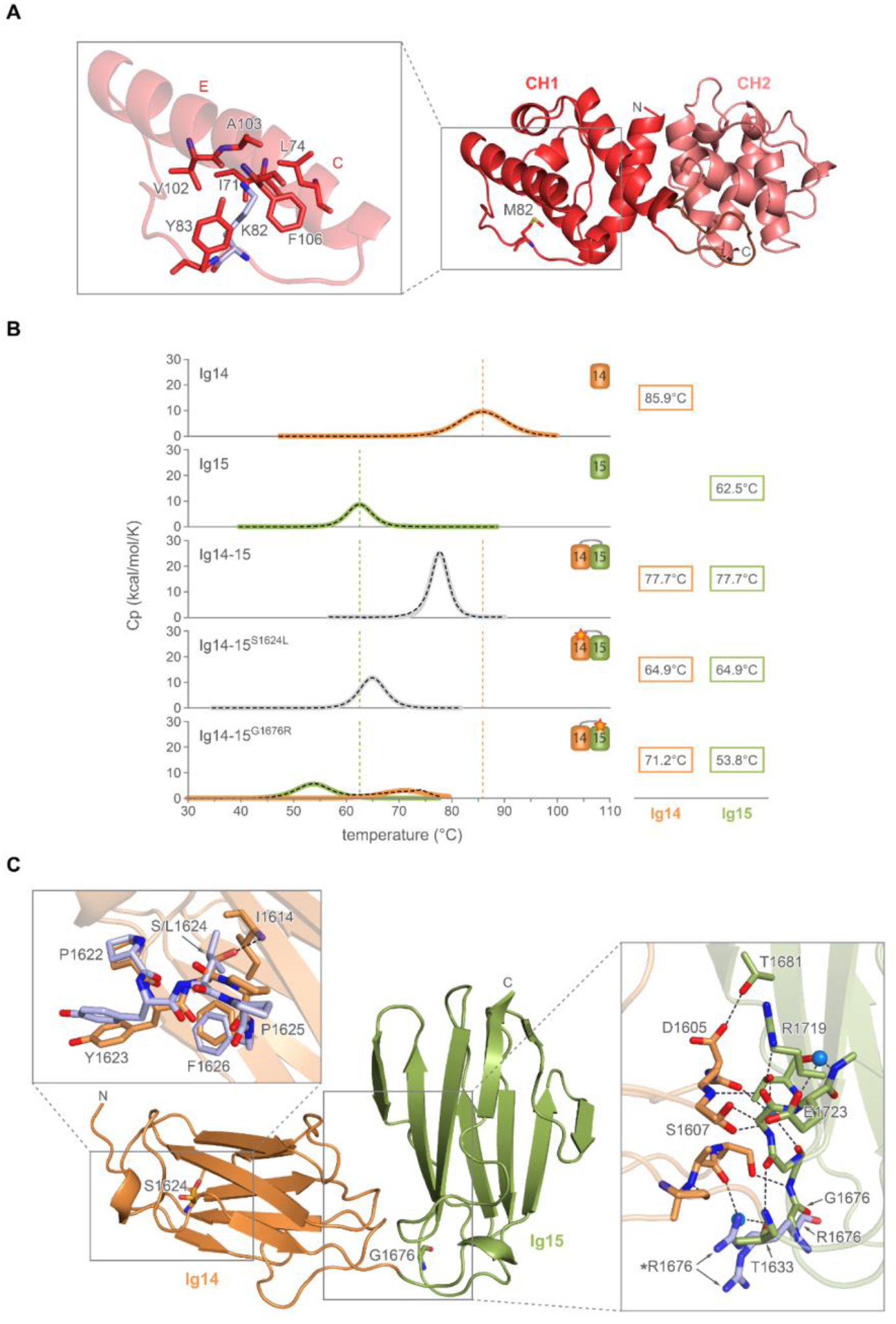
Structural and thermal stability analysis of selected mutations in ABD, and Ig14-15 domain. (**A**) ABD of human FlnC is shown in ribbon diagram, with CH1 and CH2 colored in dark and light red. Inset shows the model of p.Met82Lys mutant generated by Duet (27), with positively charged Lys side-chain clashing with the hydrophobic cluster formed by residues of h3 and h4. (**B**) Thermograms of Ig14 wild-type, Ig15wt, Ig14-15 wt, Ig14-15^G1676R^, Ig14-15^S1624L^. (**C**) Crystal structure of Ig14-15. Left inset: superposition of PXSP loops of Ig14-15^S1624L^ (blue) and Ig14-15 wild-type, showing the crucial hydrogen bond between Ser1624 to Ile1614, which is lost upon mutation to Ile, with concomitant distortion of the loop to accommodate for a bigger, hydrophobic residue at this position. Right inset: the details of hydrogen-bonding network between Ig14 and Ig15 domains, with hydrogen bonds highlighted with dashed lines, solvent molecules are shown as light blue spheres, the position of the Gly1676, mutated into Arg is labelled.

The thermal unfolding of Ig14-15 in the range 20 - 110°C showed a single transition with melting temperature (T_m_) of 77°C (**Figure 3B**). The single unfolding transition showed a ratio between Van’t Hoff and calorimetric enthalpy of 1.9, with 2.0 being expected for two equivalent domains unfolding in a single transition, thereby supporting the single entity unfolding event. The crystal structure of Ig14-15 explains and corroborates the coupled unfolding as it shows presence of extensive interdomain interactions and the formation of a hydrogen-bond stabilized interface, resulting in a single structural and folding entity (**Figures 3B and 3C**). The thermal stability of constructs Ig14-15^S1624L^ and Ig14-15^G1676R^ bearing mutations p.Ser1624Leu and p.Gly1676Arg found in patients with RCM (24), and this study), were examined under the same experimental conditions. The p.Ser1624Leu mutation is located in Ig14 (**Figure 3C<u>, left inset</u>**), and maps to a highly conserved and functionally important PXSP motif, which is present in nearly all Ig domains of human FlnC and was shown to be highly phosphorylated *in vivo* in several FlnC Ig domains (25,26). Like the wild-type, Ig14-15^S1624L^ displayed a single transition upon unfolding (**Figure 3B**), and a ratio between Van’t Hoff and calorimetric enthalpy of 1.85, in agreement with both domains unfolding in a coupled manner. In terms of fold stability Ig14-15^S1624L^ displayed T_m_ of 64.9°C, suggesting a destabilizing effect of the mutation. Comparative structural analysis showed that while the overall structure and the interaction interface between Ig14 and Ig15 appear similar, the hydrogen bond that Ser1624 side chain -OH is involved in is lost upon mutation resulting in destabilization of the interdomain interface and the decrease in T_m_ (**Figure 3C, <u>left inset</u>**). This may also explain the cellular phenotype we observed, whereby destabilization of Ig14-Ig15 interface, together with the removal of the phosphorylation site leads to the pronounced formation of Z-disc streaming in cardiomyocytes (**Figure 2**). The second mutation in this region, the p.Gly1676Arg, led to complete decoupling of the thermal unfolding of the two domains, as we observed two unfolding transitions, with T_m_s for Ig14 and Ig15 of 71.2°C and 53.8°C, respectively. The X-ray crystal structure of the Ig14-15^G1676R^ construct showed that the introduction of a larger and positively charged residue considerably perturbs the interface structure between Ig14 and Ig15, as the number of interdomain interactions mediated by water molecules increased (**Figure 3C**; **<u>Figure S4</u>**).

Based on these results, we conclude that the molecular basis of the pathogenicity of both p.Ser1624Leu and p.Gly1676Arg resides in their effect on decreasing the fold stability and/or association strength and communication between the two domains. This is further supported by our cell-based studies where we observed reduction of the cross-striated cardiomyocyte area, accompanied by the formation of extensive non-striated regions of the cytoskeleton in cardiomyocytes transfected with these variants (**Figure 2**).

We also performed analysis of thermal stability of variants where mutations are located within Ig19-Ig20 region. In addition to pathogenic mutations (p.Ile2160Phe, p.Trp2164Cys and p.Pro2298Ser), these constructs allowed us to examine two non-pathogenic ones (p.Lys2260Arg and p.Gln2058Arg). We used Ig19-Ig21 construct and observed presence of three overlapping transitions, at 60.9°C, 75.2°C and 84.8°C for the wild-type construct (**Figure 4A**). These were assigned to Ig21, Ig20, and Ig19, respectively, by comparison with the thermograms of individual domains (**<u>Figure S2</u>**). Two mutations (p.Gln2058Arg, Ig19 and p.Lys2260Arg, Ig20) displayed no impact on fold integrity of Ig19-21 constructs, in agreement with patient data as well as biophysical predictions given the relatively conservative nature of the two mutations. On the other hand, three pathogenic mutations (p.Ile2160Phe, p.Trp2164Cys, and p.Pro2298Ser, Ig20) resulted in increased stability of Ig20 (**<u>Table S1</u>**).

**Figure 4.**
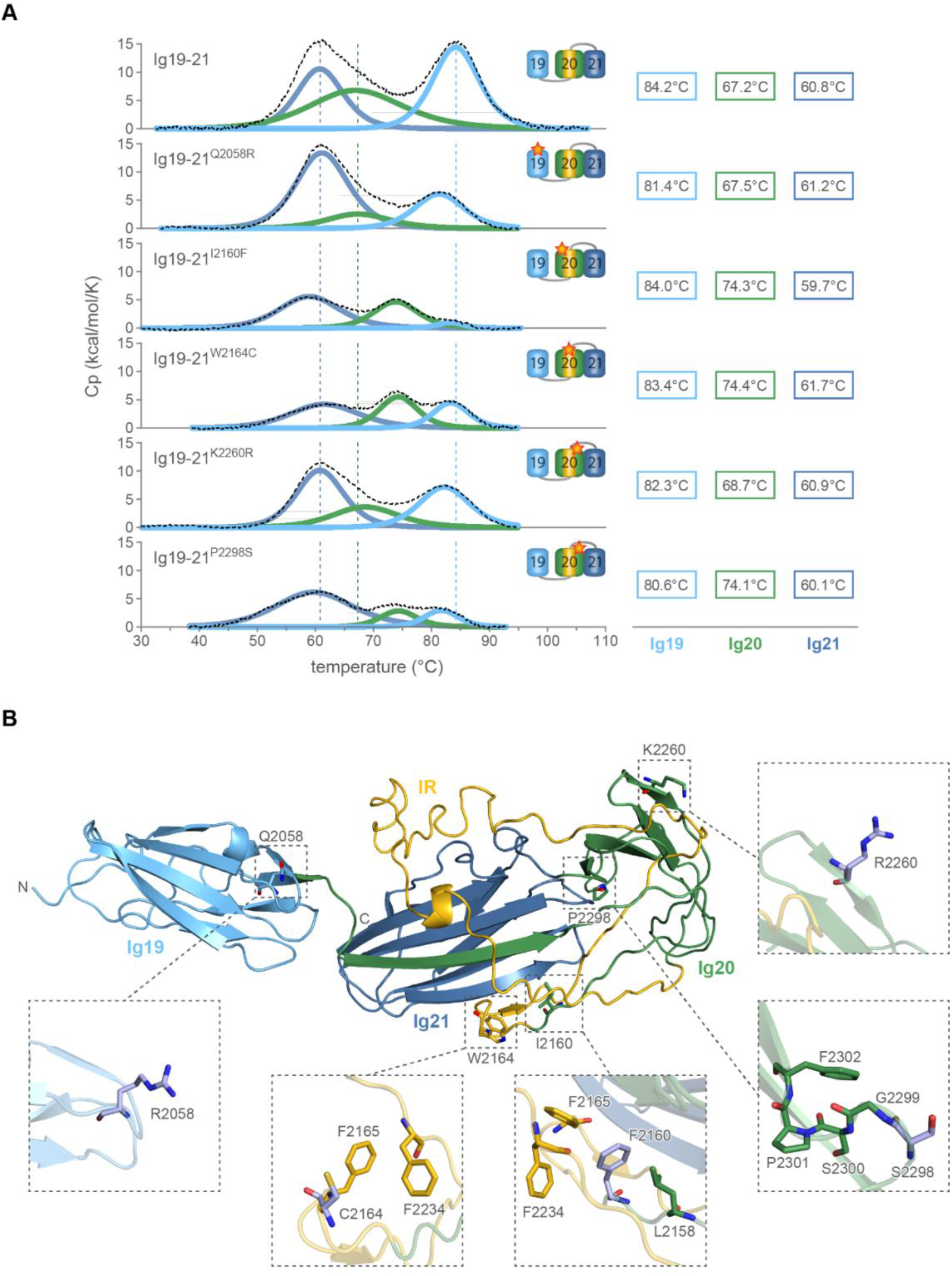
Structural and thermal stability analysis of selected mutations in Ig19-21 domains. (**A**) Thermograms of wild-type Ig19-21 and its mutants; (**B**) Cartoon diagram of Ig19-21 structural model generated by threading approach as encoded in I-tasser (30,31), with domain color code as in **A**. Positions of mutated residues p.Gln2058Arg, p.Ile2160Phe, p.Pro2298Ser, p.Lys2260Arg, p.Trp2164Cys are depicted as stick models. A detailed view of the mutations generated with Duet (27) are shown in the insets.

Mutation p.Ile2160Phe, associated with RCM (24), closely precedes the IR in Ig20 (aa 2162-2243) (**Figure 4B, <u>Figure S3</u>**), and leads to stabilization of Ig20 compared to the wild-type by 7°C, while the other domains are not affected. The stabilization of the mutant Ig20 is due to aromatic stacking interactions with other aromatic residues that Phe2160 can establish (**Figure 4B**). A close by variant p.Trp2164Cys, associated with HCM (H. Watkins, personal communication), is particularly interesting since it maps to the beginning of the IR which is considered to lack a unique and well-defined 3D structure. The mutation leads to an increase in the melting temperature of Ig20 by 7°C, suggesting a local structuring rearrangement, as inferred from the relative increase in enthalpy change upon unfolding of Ig20 compared to Ig19 and Ig21 (**Figure 4A**). In the case of p.Ile2160Phe and p.Trp2164Cys, we hypothesise that increased relative enthalpy of unfolding, suggestive of an increased structuring of the surrounding of the mutation, can negatively impact interactions with binding partners, where structural plasticity of a disordered region is required, leading to RCM (24) or HCM (H. Watkins, personal communication), respectively.

Additionally, p.Pro2298Ser mutation, associated with RCM (this study and (28)) maps to Ig21 (**Figure 4B**), and shares an important similarity with p.Ser1624Leu in Ig14: both mutations are part of the highly conserved PXSP motif (23). Similar to the p.Ser1624Leu, the p.Pro 2298Ser mutation also influences the stability of neighbouring domains (**Figure 4B**), possibly through inter-domain contacts that have been reported previously (18,19,29) Ruskamo, 2012 #133}. The observed increased T_m_ (7°C) could be attributed to the replacement of Pro with Ser, which can form additional hydrogen bonds, both with the side-chain and the main-chain groups. The mutation p.Pro2298Ser affects the first proline residue in the conserved PXSP motif, which was shown to be involved in the regulation of mechanosensing in FlnA Ig20 through *cis-trans* isomerisation of the second proline of the motif (23). Substitution of proline with a multivalent hydrogen-bond acceptor and donor residue, serine, could therefore disturb the finely tuned *cis-trans* isomerisation balance involved in mechanosensing regulation.

Taken together, our biophysical and structural data provide a basis for molecular understanding of the effect point mutations have. We observed that some pathogenic mutations destabilize while others stabilize the FlnC variants. At the same time, p.Gln2058Arg and p.Lys2260Arg mutations have no effect on stability, suggesting that either destabilization or stabilization of FlnC can have a negative impact on its ability to function properly, resulting in pathogenesis.

### Pathogenic variant mapping and grouping in a representative Ig domain

Finally, we examined whether the location of pathogenic mutations matches with biophysical and structural expectations. To address this, we overlayed structures of all 24 Ig domains (either experimental or predicted) with the structure of Ig5, which was selected as the reference since it shows the lowest average RMSD with other Ig domains (RMSD = 0.64 Å -2.467 Å, with an average value of 1.15 Å). The pathogenic mutation sites were mapped on the structure-based sequence alignment derived from the above superpositions. We observed that mutations were not randomly distributed, but clustered in a few regions (**Figure 5A)**, and that the frequently mutated sites are conserved in FlnC Ig domains (**Figure 5B)**. This is in good agreement with widely accepted view that residues critical for folding and/or function display higher levels of conservation within a domain family (such as Ig family analyzed here), and that mutations at these sites are more likely to be detrimental.

**Figure 5.**
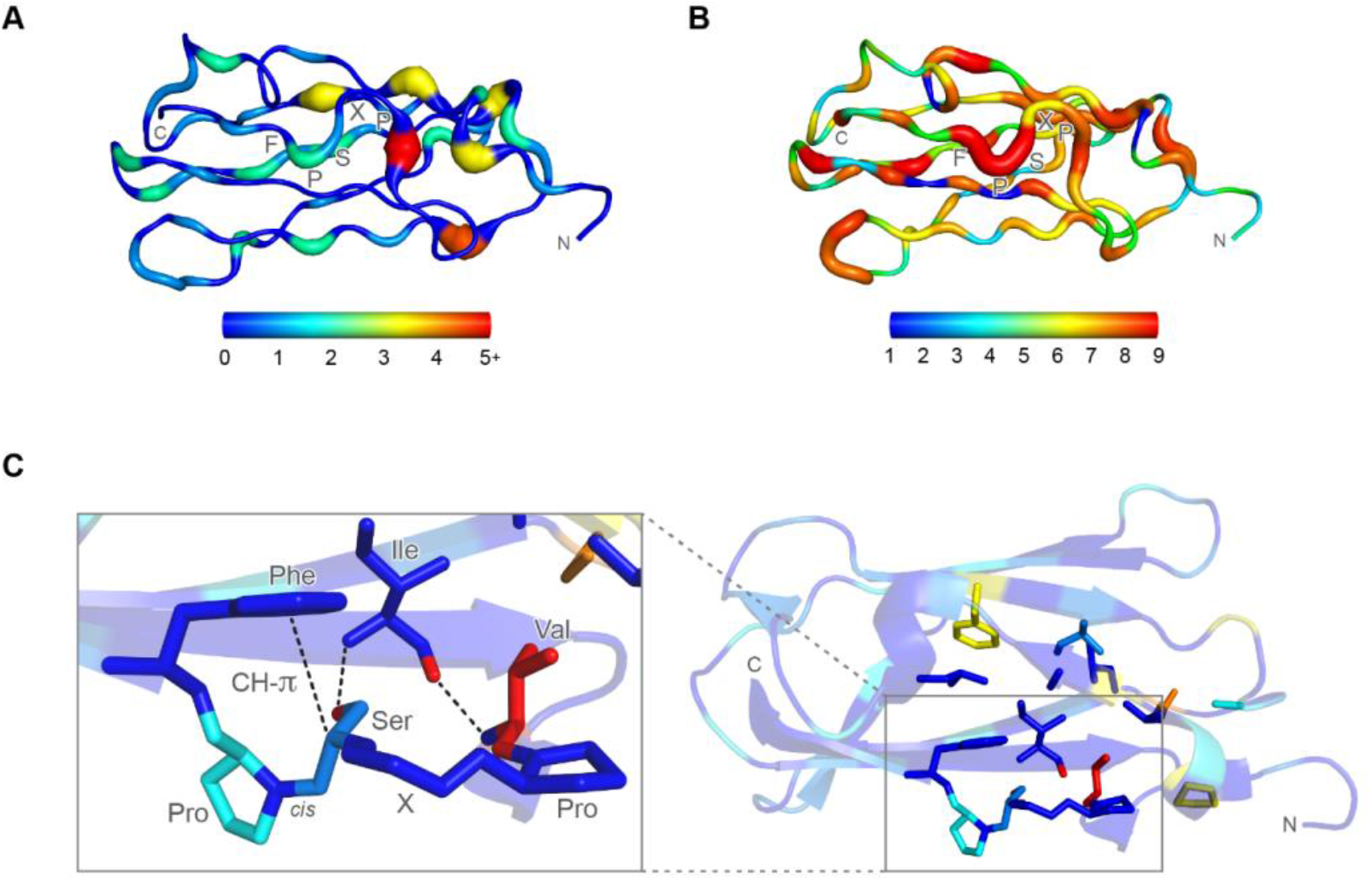
Frequency of pathogenic mutations and amino acid conservation in human FlnC Ig domains mapped on the representative structure of Ig5. (A) Frequency of pathogenic mutations. Positions that do not host pathogenic mutations are shown in blue and with a narrow ribbon. Positions that host five (or plus) pathogenic mutations are shown in a red and thick ribbon. Residues with intermediate numbers of pathogenic mutations are represented according to the color code shown in the bar shown below; (B) Degree of conservation of residues in human FlnC Ig domains. The conservation score, determined by ConSurf (32) is mapped according to the bar shown below, with blue and narrow ribbon corresponding to low, and red and large ribbon corresponding to high conservation. PXSP motif is labeled in both panels; (C) PXSP loop and surrounding hydrophobic cluster, colored according to the mutational frequency as shown in A. Inset: The crucial hydrogen-bond between the serine side-chain oxygen and Leu main-chain nitrogen residing on the adjacent β-strand is shown. A highly conserved hydrophobic residue colored in red (either valine or isoleucine in all Ig domains) forms another hydrogen main-chain main-chain hydrogen bond with Leu, further stabilising the structure. Carbonyl oxygens omitted with the exception of the relevant hydrogen bond engaging one for clarity.

### Development of predictive bioinformatics tool AMIVA-F: the training set

Since a growing number of FLNC variants are associated with (cardio)myopathies, we were motivated to develop a prediction tool that uses machine learning (Analysis of Missense Variants in human Filamin C, AMIVA-F) to predict variants’ pathogenicity. To develop such predictive bioinformatic tool, we first selected a training set of variants associated with HCM, DCM, RCM, myofibrillar and distal myopathy, and several others. A total of over 250 unique variants were retrieved from the international peer-reviewed literature. Variants that lacked a clear description of the associated phenotype or clinical information were not included in the collection. Furthermore, we only collected single point missense variants, excluding frameshift or truncating variants, resulting in a total of 108 disease related variants (**Figure 6**). Each variant in the disease-related dataset was associate with pathogenicity in at least one peer reviewed publication (**<u>Table S3</u>**). Out of 108 variants we chose, 65 variants are associated with HCM, eight with RCM, five with myofibrillar myopathies, five with distal myopathies, four with ACM, three with DCM, two with IBM, two with congenital heart disease, two with left ventricular non-compaction, and 12 with other disease phenotypes.

**Figure 6.**
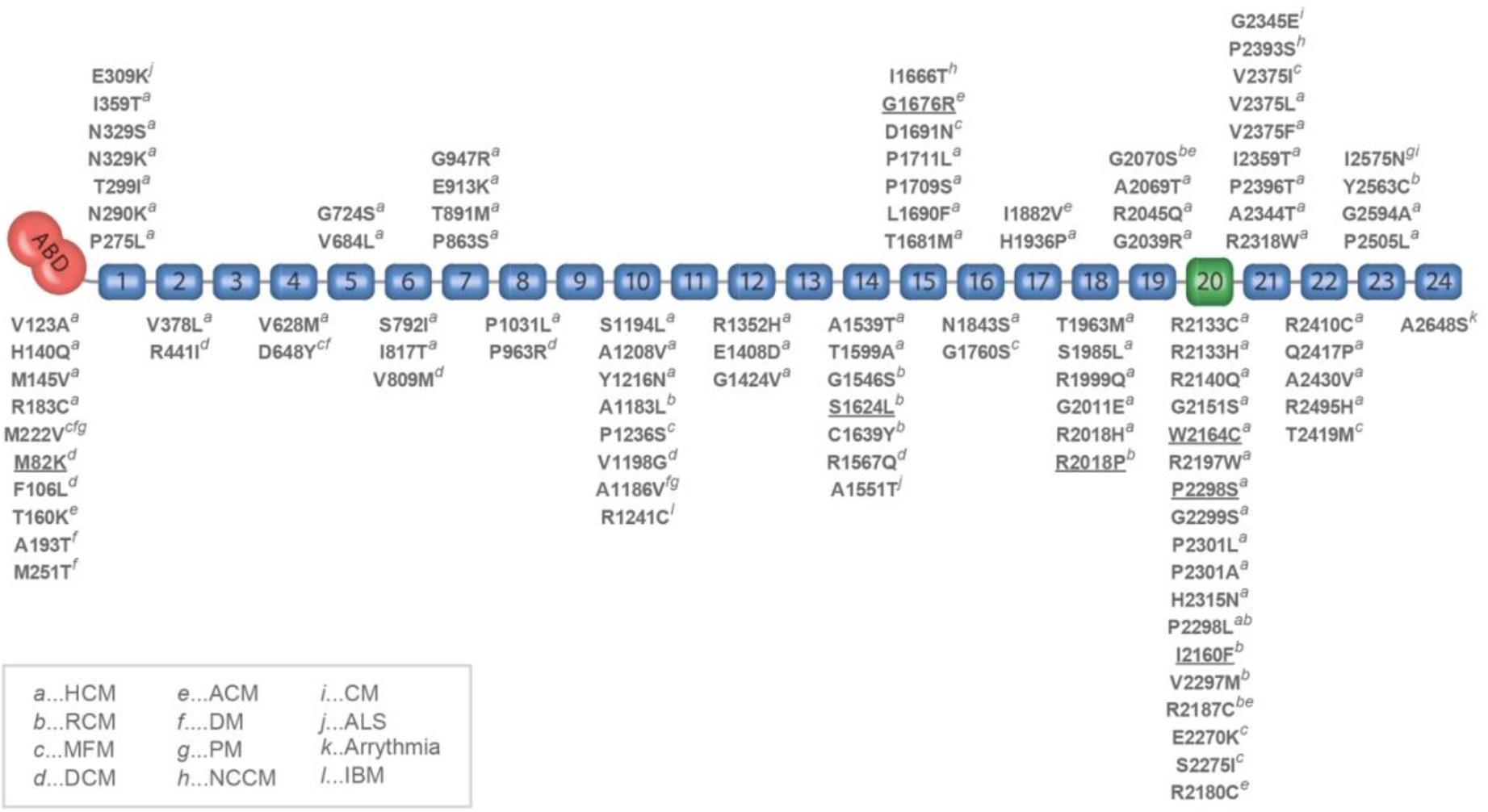
Overview of *FLNC* variants. Variants in the *FLNC* gene that have been previously or in this study reported in individuals with myopathy or cardiomyopathy, mapped to the corresponding FlnC domain. Variants are annotated at protein level. Abbreviations: HCM (hypertrophic cardiomyopathy), RCM (restrictive cardiomyopathy), MFM (myofibrillar myopathy), DCM (dilated cardiomyopathy), ACM (arrhythmogenic cardiomyopathy), DM (distal myopathy), PM (peripheral myopathy), NCCM (non-compaction cardiomyopathy), CM (cardiomyopathy), ALS (amyotrophic lateral sclerosis), IBM (inclusion body myositis). Mutants experimentally characterized in this work are underlined.

For the selection of a negative or neutral mutation set, we gathered single point missense variants found in the Genome Aggregation Database (gnomAD; https://gnomad.broadinstitute.org). This yielded a collection of 1500 missense variants, which were analyzed and selected according to the following criteria: (i) they were found at higher frequency, as recent study suggested that loss of function and disease related mutations tend to be found at extremely low frequencies compared to neutral ones (33); and (ii) the presence of individuals having homozygous alleles for variants was taken as further evidence of a variant being neutral. We selected variants with high allele frequency (> 10^-3^), and high allele counts based on these criteria. By comparison, variants found in the gnomAD associated with pathogenicity were found at frequencies around 10^-6^, a difference of up to three orders of magnitude. We discarded variants that showed less than 10 allele counts as these variants might be rare but harmless. In total, these stringent criteria selections led to a reduction from 1500 to 65 variants and constituted the neutral dataset. The average allele count in the neutral variant dataset was 504, and some selected variants also showed homozygous alleles.

In summary, our two datasets, consisting of 108 disease-related variants (**Figure 2, <u>Table S3</u>**) and 65 presumably neutral variants (**<u>Table S4</u>**), were merged to 173 instances and used as training sets for our algorithm. An overview of the distribution of pathogenic variants through FlnC domains is reported in **Figure 6**, showing that they tend to cluster in the C-terminal region (**Figure 1)**, which is a mutational and protein-protein interaction hot-spot of FlnC (4,8,11,34). The training set allowed us to observe that mutations of arginines and methionines result in pathogenicity more frequently than expected based on amino-acid composition, in particular when arginine is mutated into cysteine. Pertinent details on the propensity of each amino acid type to result in pathogenic variants upon mutation are reported in the **Supplementary Information.**

### Defining the attributes for AMIVA-F analysis

AMIVA-F tool is based on the hypothesis that pathogenic variants modify protein function in a complex way by impacting the fold stability and integrity as well as by influencing interaction interfaces or phosphorylation sites. To characterize the impact of variants at a biophysical and structural level, we first prepared a collection of 3D structures for all of wild-type FlnC domains that included 6 experimental structures covering 8 Ig domains, and 15 3D computational models covering the ABD and 16 Ig domains (**<u>Figure S1</u>, <u>Table S5</u>**).

In order to fully characterize a variant, we defined 11 different attributes (see **Materials and Methods** for details). These attributes were derived through analysis of 3D structures by using scripts included in the AMIVA-F package. Several attributes are related to the solvent accessibility of the residues: the absolute solvent accessible surface areas of both wild-type and mutant residues, and the discretized solvent accessibilities, grouped into the three categories (inaccessible, partially accessible, and accessible, as defined by Worth et al., (2011) (35), and detailed in **Materials and Methods**). Other attributes are related to residue properties, like the change in hydrophobicity upon mutation, according to the scale defined by Kyte et al. (1982) (36), the spatial aggregation propensity score (SAP-score) (37), the change in the number of non-hydrogen atoms upon mutation, and the secondary structure of the residue that is mutated. Furthermore, we included attributes that do not depend just on residue properties: the side-chain orientation of the mutated residue - this is ignored for Gly and Pro residues - and a composite variable, taking into account known binding partners, side-chain clashes introduced upon mutation as well as potential disruption of phosphorylation sites.

### Benchmarking and validating AMIVA-F

Predictions were made with a multilayer perceptron neural network, by using WEKA (38), an open-source workbench that allows the use of several machine learning techniques. In order to benchmark the performance of AMIVA-F, we used ZeroR algorithm (38) that predicts the most common instance in a dataset and therefore defines a baseline accuracy. In general, baseline accuracy is solely dependent on the underlying dataset, and it is algorithm independent. For our dataset, ZeroR determined 62.4% as baseline accuracy.

Given the relatively low number of instances (173 variants), particular precautions had to be taken to train the network because, upon regular cross-validation, there is a probability to bin outliers together and artificially increase bias. For larger datasets, this concern becomes less relevant given that binning large datasets results in a lower contribution of potential outliers to the binned averages. In comparison, in smaller datasets, a few outliers could considerably skew the average. To counteract that, we sampled by using 10-fold stratified cross-validation to ensure equal incorporation of all data while also preventing disproportionate bias through random sampling, which could happen in regular unstratified cross-validation. **Table 1**. Overall, sensitivity and accuracy are close to or slightly over 0.8 as well as the area under the ROC curve (AUC) and the F-statistics; specificity, though slightly lower, is still in a comfortable range close to 0.7 (see **Materials and Methods** for details).

**Table 1.** Summary of the quality of the predictions of AMIVA-F.

| Table 1<br>Summary of the quality of the predictions of AMIVA-F. |  |
| --- | --- |
| **Sensitivity**<br>$(TP^{(a)} / (TP + FN^{(b)}))$ | 0.858 |
| **Specificity**<br>$(TN^{(c)} / (TN + FP^{(d)}))$ | 0.689 |
| **Accuracy**<br>$(TP+TN)/(TP+TN+FP+FN)$ | 0.786 |
| **Precision**<br>$(TP / (TP + FP))$ | 0.795 |
| **F-measure**<br>$TP / (TP + \frac{1}{2} * (FP + FN))$ | 0.788 |
| **ROC-Area under the curve (AUC)<sup>(e)</sup>** | 0.818 |
| **MCC<sup>(f)</sup>**<br>$(TP*TN)(PF*FN)/[(TN+FN)(TN+FP)(TP+FN)(TP+FP)]^{1/2}$ | 0.560 |

To validate the AMIVA-F performance further, we used 8 variants classified to be likely pathogenic according to criteria of ACMG and Fokkema et al (2011) (40) as external test cases. Their exclusion from the training sets was justified by the absence of concrete information about the specific disease underlying those variants, even though the probability of the variant to be disease causing is sufficient to warrant clinical actions (41). As shown in **Table 2**, six predictions were correct (75%), in line with the cross-validation of AMIVA-F and this reinforces the cross-validation estimations, despite the small dimension of the external test set (8 variants).

**Table 2.**
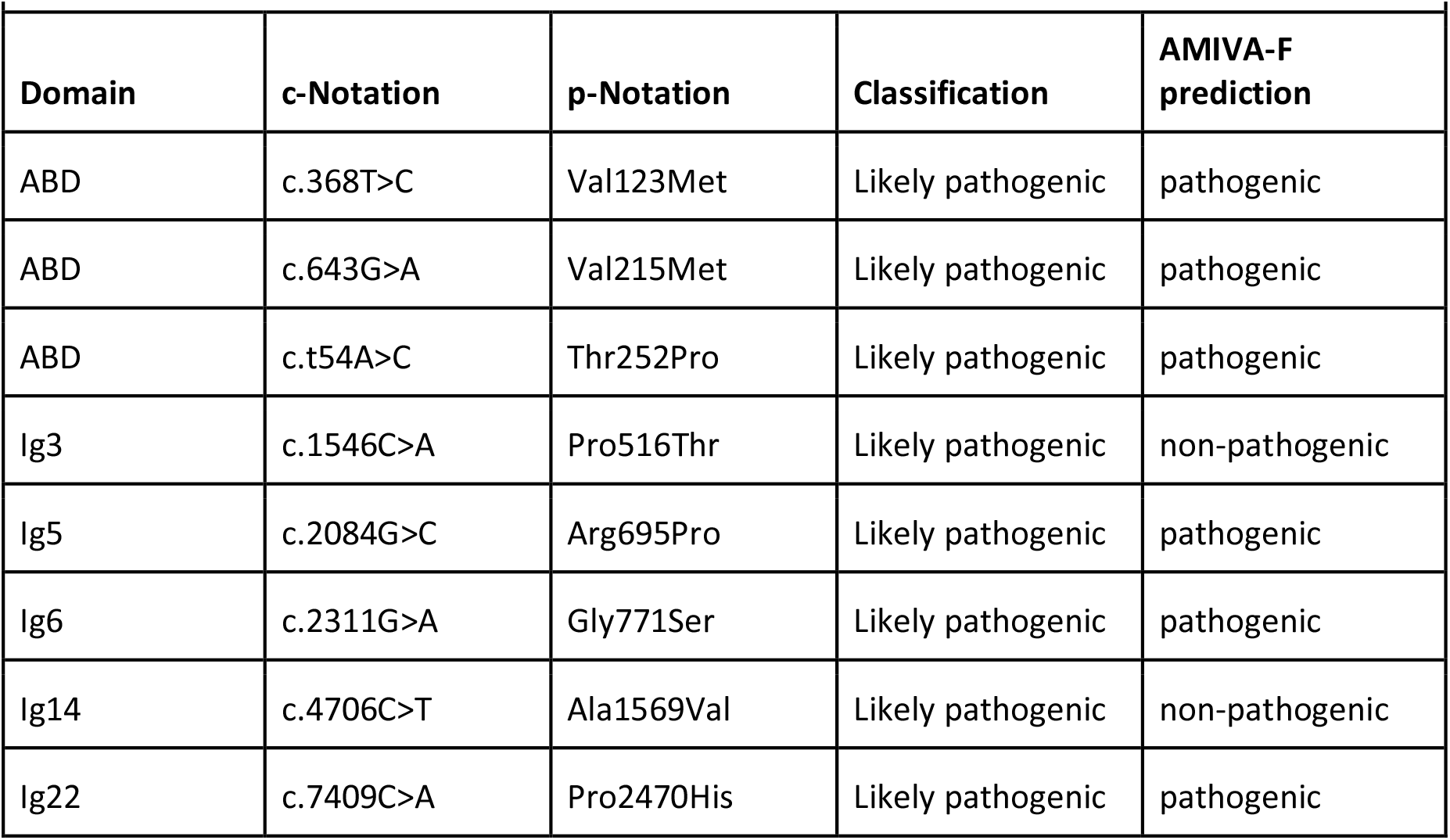
Pathogenicity prediction of eight “likely pathogenic” classified variants which were absent from the training set.

|  |  |  |  |  |
| --- | --- | --- | --- | --- |
| Domain | c-Notation | p-Notation | Classification | AMIVA-F prediction |
| ABD | c.368T>C | Val123Met | Likely pathogenic | pathogenic |
| ABD | c.643G>A | Val215Met | Likely pathogenic | pathogenic |
| ABD | c.t54A>C | Thr252Pro | Likely pathogenic | pathogenic |
| Ig3 | c.1546C>A | Pro516Thr | Likely pathogenic | non-pathogenic |
| Ig5 | c.2084G>C | Arg695Pro | Likely pathogenic | pathogenic |
| Ig6 | c.2311G>A | Gly771Ser | Likely pathogenic | pathogenic |
| Ig14 | c.4706C>T | Ala1569Val | Likely pathogenic | non-pathogenic |
| Ig22 | c.7409C>A | Pro2470His | Likely pathogenic | pathogenic |

Additionally, we examined whether predictions are biased by the frequency of occurrence of the variants in the GnomAD database. This was an important point to check because our non-pathogenic variant learning set was assembled based on the premise that non-pathogenic variants exhibit high occurrence frequency (33,42). Furthermore, all variants that we included in the pathogenic variant learning are infrequent variants, although their frequency was not part of selection criteria. For this purpose, 16 rare variants (<10^-6^ allele frequency), with no reported pathogenicity, were selected and used as test cases. Nine of them were predicted to be pathogenic and seven to be non-pathogenic (**<u>Table S6</u>**). While there is no experimental or observational proof that these predictions are correct, they indicate that the algorithm is not simply classifying based on rarity. In other words, this indicates that predictions made with AMIVA-F are independent of the variant frequency within the population. Pathogenic variants identified by AMIVA-F match well with biophysical and structural expectations regarding their effects. Collectively, our benchmarking and validation indicate that AMIVA-F performance is adequate to make accurate and actionable predictions.

## Discussion

Many patients suffering from cardiomyopathies carry mutations in the *FLNC* gene. Therefore, early identification of disease-associated mutations in this gene is becoming particularly important from a clinical perspective. However, research has shown that not all *FLNC* mutations are pathogenic, and some cardiomyopathies are caused by other factors, highlighting the need for methods that can predict pathogenicity of *FLNC* mutations. In this context, knowledge-based machine learning algorithms may represent a powerful strategy for rapid *FLNC* variant classification. Here, we address this need by combining clinical information with cellular, molecular and biophysical data to develop computational tool for automatic identification of pathogenic FlnC variants. The unique strength of our approach is the seamless integration of patient data, which resulted in description of eight *FLNC* variants, and cell-based, biophysical and structural analysis, which characterized the effect of several variants with varied disease severity (from neutral to highly pathogenic). These data, together with comprehensive information gathered from the literature, yielded the machine-learning algorithm AMIVA-F that predicts pathogenicity of FlnC variants with actionable accuracy.

In terms of our patient data, this study investigated the following clinical phenotypes: (i) the most severe phenotype DCM/ARVC with heart failure resulting in HTX at 3rd decade of life (patient #1); (ii) two female patients (patient #6 and #7) with mild cardiac abnormalities who did not develop CM phenotype until the 8th decade of life, and had a history of SD in the family due to ARVC diagnosed PM; and (iii) RCM/HCM phenotype, also overlapping in the families with dominant atrial tachyarrhythmias and HF presentation, no sustained VT, no sudden death in the family, and without peripheral myopathy (patients #2 - #5, #8). In addition to differences in clinical phenotypes, we showed that patients in our cohort also exhibited differences in *FLNC* variants. The most sever phenotype (patient #1) was the carrier of variant c.245T.G that results in p.Met82Lys point mutation located within the ABD of FlnC, which is likely related to protein aggregate toxicity, similarly to myofibillar myopathy or HCM (43) (5). Thus, this is in general agreement with a recognized role of *FLNC* variants that cause misfolding in the mechanism of CMs (3). Of the other patients in our cohort, mutations found in three patients (p.Ser1624Leu in Ig14 (Patients #2 and #3); and p.Lys2260Arg in Ig20 (Patient #7)) were classified as “conflicting interpretation of pathogenicity” by ACMG. Additionally, two mutations (p.Gly1676Arg in Ig15 (Patient #4) and p.Gln2058Arg in Ig19 (Patient #6)) are classified as being of “uncertain significance” by ACMG, whereas two mutations (p.Arg2018Pro in Ig18 (Patient #5) and p.Pro2298Leu in Ig20 (Patient #8) are not currently classified by ACMG. The lack of classification or uncertainty and conflicting interpretation of pathogenicity all contribute to current difficulties in CM patient stratification and treatment.

Our results are of special interest and significance given that majority of best characterized disease-associated *FLNC* variants are truncations, or changes with ABD and ROD 1 and ROD 2 domains (6,44–46). Only three missense FLNC variants have been classified as likely pathogenic: p.Phe106Leu in the setting of compound heterozygozity, p.Ala123Met, and p.Gly2070Ser, the remaining were classified as variant of uncertain significance (VUS) (4). This emphasises the requirement for a more refined categorization of the *FLNC* missense variants. Overall, although the majority of the presented variants are VUS according to ACMG, we provided a compelling clinical, genetic and experimental evidence of the involvement of 6 of 8 missense *FLNC* variants in CM, with the final classification of gene variants based on the bioinformatics tool AMIVA-F and familial cosegregation studies.

Our systematic analysis of the effects of point mutations on stability, structure and function of FlnC yielded several important insights. As noted before, pathogenic point mutations are distributed across the entire FlnC protein - across its ABD as well as the 24 Ig domains. Therefore, we selected to conduct an in-depth characterization of point mutations distributed across different domains. In our hands, the construct featuring a point mutation within the ABD (Met82Lys) was not stable enough to be recombinantly expressed. This mutation also caused a profound phenotype in our cell-based assays, where we observed formation of cytoplasmic FlnC-containing aggregates that sequestered titin, leading to disappearance of striated myofibrils. Two pathogenic mutations were analyzed, p.Ser1624Leu and p.Gly1676Arg, which we studied in the context of the Ig14-15 construct, decreased thermal stability and disrupted its structure. Additionally, both p.Ser1624Leu and p.Gly1676Arg also had a profound phenotypic effect in cells, in further agreement with severity of the disease documented in patients. Therefore, point mutations that destabilize FlnC have a negative impact on its function, possibly by disrupting its proper folding necessary for establishing correct interactions with numerous functionally relevant binding partners. This is in agreement with commonly accepted view that mutations that destabilize protein structure are detrimental to their function, as well as our results obtained using variants (p.Gln2058Arg in Ig19 and p.Lys2260Arg in Ig20) that exhibited no effect on thermal stability or structure, suggesting that they might be non-pathogenic. Unexpectedly, pathogenic mutations (p.Ile2160Phe and p.Pro2298Ser studied in the context of Ig19-Ig21 construct), had the opposite effect on thermal stability. These mutants were more stable than the wild-type, and yet were associated with severe disease phenotypes. This suggests that stabilizing changes in FlnC structure can also have a negative impact on its ability to engage relevant binding partners, thus interfering with its function. Overall, our results indicate that the relationship between variant pathogenicity and thermal stability of the human FlnC domains is rather complex and can be unexpected. However, two conclusions are clearly supported by our data: (i) pathogenicity is associated with perturbations of fold stability; and (ii) pathogenicity can be associated with inter-domain interactions, suggesting that biophysical pathogenicity-stability studies need to be conducted not with single but multi-domain constructs. Furthermore, when we overlaid structures (solved or calculated) of all Ig domains in FlnC and mapped distribution of pathogenic mutations in our dataset, we observed that these cluster within specific structural elements, most notably between the loop hosting the PXSP motif and the N-terminal side of the Ig domain (**Figure 5A**). These regions are highly conserved among human FlnC Ig domains (**Figure 5B, Figure S3**), suggesting that structural conservation combined with sequence conservation might be a stronger indicator of functional relevance, and, by extension, predictor of pathogenicity.

Driven by our interest to understand molecular basis of pathogenic mutations, we examined the interactions of PXSP motif and noted a presence of phenylalanine following the PXSP motif. This residue forms CH-π electron interactions that can stabilize the uncommon Pro *cis* conformation (47) (**Figure 5C<u>, inlet</u>**), and contacts additional regions including a highly conserved Ile/Val residue that precedes the motif and is a pathogenic hotspot (**Figure 5C**). Given the importance of the phenylalanine residue in orchestrating both the PXSP motif and the stability of the Ig domain, we propose extending the motif to PXSPF. Further evidence for the critical role of the PXSPF motif and specifically of the serine residue comes from a phosphoproteomics study (26), which revealed this site is phosphorylated in several Ig domains of FlnC. Our data corroborate its importance, as mutation of this serine is one among several pathogenic mutations in the PXSPF motif. Based on its high conservation, we hypothesize that mutations of the PXSPF motif in other domains will likely be pathogenic as well. Additionally, we also note the presence of pathogenic mutations with a unique 82 amino acid insertion specific to FlnC (when compared to other family members).

We used what we learned from our studies and combined it with literature information to develop a pathogenicity prediction machine-learning algorithm AMIVA-F based on a multilayer perceptron neural network. Its performance, estimated by 10-fold stratified cross-validation and using an external test set, is remarkable as AMIVA-F outperforms other predictive tools, such as Polyphen2 (48), SIFT (49) and Provean (50), used in clinical diagnostics designed to predict changes in protein stability (28). AMIVA-F increases the accuracy to 78.6% against 73.4%, 64.2%, and 65.9% for Polyphen2, SIFT, and Provean, respectively (**<u>Table S7</u>**). However, Polyphen2, SIFT, and Provean are applicable to all types of proteins, while AMIVA-F was specifically designed to analyze variants in FlnC. This suggests that using protein-specific knowledge, such as structural, biochemical, and biophysical attributes, improves prediction accuracy, and may therefore represent an important approach going forward. If structural data and suitably large learning sets are available, similar machine learning techniques can be tailored to analyze missense variants of any given system.

## Materials and methods

### Patient data

The patients described in the paper were recruited to a multicenter, observational, longitudinal cohort study of patients with FlnC recruited from 19 European cardiomyopathy units between 1990 and 2018. The study conforms to the principles of the Helsinki declaration. The authors from each participating center guarantee the integrity of data from their institution and have approval for anonymised patient data collation and analysis from local ethics committee/internal review board. All participating patients consented to genetic testing. Detailed methods on the genetic testing and clinical assessment were published recently (45).

### Cell biophysics

Neonatal rat cardiomyocytes were prepared by standard methods from 3-day old pubs, plated on collagen-coated dishes and transfected one day after plating with pEGFP constructs containing wild-type or mutant filamin-C, tagged at the N-terminus with GFP, as described (gift of D. Fürst, Bonn. (13,51). After two days, cells were fixed with 4% paraformaldehyde on ice for 5 minutes, processed for immunostaining with the T12 anti-titin monoclonal antibody (52) and imaged on a Zeiss LSM510 Meta confocal microscope using a 63X oil immersion objective.

### Structures and structural models of FlnC Ig-domains

The 3D structures of 25 filamin domains that have been determined experimentally were downloaded from the Protein Data Bank (53–55) (**<u>Table S5</u>**). For domains where experimental 3D structures were unavailable, homology models were generated by using the Modeller package (56), based on available structures of FlnA or FlnB templates. A total of five structures per template were generated based on Modeller Automodel function, energy was minimized according to DOPE and GA341 scores and the lowest energy model was used for further computations (systematically associated with a GA341 score of 1.00). For Ig16 and Ig19-21, we utilized I-Tasser (30,31) models which were generated through threading. In order to reinforce our Modeller and I-Tasser homology structures, we used AlphaFold2 (57) which showed excellent agreement. Ribbon representations of all structures can be found in **<u>Figure S1</u>**.

### Training variants learning sets

In order to distinguish between pathogenic and benign variants of FlnC mutations we gathered a total of 173 mutations and split them into two classes.

The first class consisted of disease-related mutations which were already verified experimentally and were classified as “pathogenic” (**<u>Table S3</u>**).

For the second class of mutations, we selected missense mutations from the GnomAD database according to the criteria suggested by Karczewski et al. (33), who found that pathogenic mutations tend to appear with lower frequencies, compared to neutral ones. Furthermore, the presence of homozygotes can give indication that a mutation is potentially benign. In detail, non-pathogenic variants were selected according to the following criteria: (i) high allele count (number of cases reported), (ii) allele frequency >> 10^-3^ (extrapolation of frequency in the population), and/or (iii) presence of homozygotes. Details on the selection of non-pathogenic variants can be found in **<u>Table</u> <u>S4</u>**.

In total, this resulted in 108 pathogenic and 65 non-pathogenic variants.

### Machine learning algorithm

We utilized WEKA (38) a workbench for machine learning tools and algorithms in order to set up our machine learning algorithm.

In order to benchmark our algorithm, we used ZeroR, an algorithm that always predicts the most common instance in a dataset and therefore defined a baseline accuracy, which is solely dependent on the underlying datasets and is algorithm independent. For our dataset, ZeroR determined a 62.4% baseline accuracy.

Initially, we tried several algorithms, ranging from random forest type algorithms, over decision tables, bayesian logic, linear regressions and many more. Ultimately, the final optimized algorithm was found to be a multilayer perceptron neural network with a weighted average true positive (TP) rate of ∼80%. The exact metrics are shown in (**Table 1**). Pertinent details are shown in **<u>Table S8</u>**.

### Cross-validation and prevention of overfitting

Due to the relatively low amount of data (173 variants), particular precautions needed to be taken. This arose due to the fact that upon normal cross validation, probability would be higher to bin outliers together and therefore artificially increase bias. For bigger datasets, this concern becomes less relevant given that binning large datasets results in lower contribution of potential outliers to the binned averages while in smaller datasets, a few outliers could skew considerably the average. In order to counteract that, we utilized 10-fold stratified cross validation to ensure incorporation of all data, while also preventing disproportionate bias through random sampling which could happen in normal unstratified cross validation.

### Attributes used to distinguish pathogenic and benign mutations

The 11 different parameters used to feed our algorithm are described below

1. Absolute solvent accessibility in Å^2^ of the wild-type residue, computed with Naccess (Hubbard, S J.,Thornton, J M. NACCESS, Department of Biochemistry and Molecular Biology, University College London)
2. Absolute solvent accessibility in Å^2^ of the mutant residue, computed with Naccess (Hubbard, S J.,Thornton, J M. NACCESS, Department of Biochemistry and Molecular Biology, University College London)
3. Relative solvent accessibility of the wild-type residue, discretized into three classes according to **(WORTH; PREISSNER; BLUNDELL, 2011)**: accessible (>43% relative SASA), partially accessible (17-43% relative SASA), and inaccessible (<17% relative SASA).
4. Relative solvent accessibility of the mutated residue, discretized into three classes according to **(WORTH; PREISSNER; BLUNDELL, 2011)**: accessible (>43% relative SASA), partially accessible (17-43% relative SASA), and inaccessible (<17% relative SASA).
5. Change in numbers of non-hydrogen atoms upon mutation.
6. Change in hydrophobicity upon mutation according to the scale of (KYTE; DOOLITTLE, 1982).
7. Difference between attributes 3 and 4, which are discrete variables: as a consequence this attribute is discretized too and it is classified at “no change” if the attributes 3 and 4 are equal, “better” if the solvent accessibility increases upon mutation, and “worse”in the opposite case.
8. A composite discrete variable, taking into account known binding partners, introduces side-chain clashes upon mutation as well as potential disruption of a phosphosite. Mutations were deemed to clash if there was no side-chain rotamer in the Pymol library (The PyMOL Molecular Graphics System, version 2.3, Schrödinger, LLC) that could have been fitted without altering the structure. Mutations were considered to be related to protein phosphorylation, and as a consequence prone to engender pathological consequences, if closer than 8 Å from a phosphorylation site. Cα - Cα - distances were used as a cutoff distance here and phosphorylation sites were taken from (REIMANN; SCHWÄBLE; FRICKE; MÜHLHÄUSER et al., 2020).
9. side-chain orientation of the affected amino acid. In the case of Pro/Gly this is labelled with a “?”. This attribute can either assume 2 states, “towards solvent” or “towards protein core”.
10. Type of secondary structure of the mutated residues - a special tag was used for mutations located in the partially disordered insertion region of Ig20.
11. A composite variable, combining solvent accessibility and change in hydrophobicity named SAP-score (VOYNOV; CHENNAMSETTY; KAYSER; HELK *et al.*, 2009). This parameter was shown to correlate well with aggregation propensity, hence the name spatial aggregation propensity score.

### Plasmids and DNA constructs

Ig14 (residues 1534-1634), Ig15 (residues 1633-1736), Ig14-15 (residues 1534-1736), Ig14-15^S1624L^(residues 1534-1736), Ig14-15^G1676R^ (residues 1534-1736), Ig19 (residues 2033-2130), Ig21 (residues 2313-2406), Ig2021 (residues 2132-2406), Ig1921 (residues 2036-2406), Ig1921^I2160F^ (residues 2036-2406), Ig1921^W2164C^ (residues 2036-2406), Ig1921^P^ ^2298S^ (residues 2036-2406), Ig1921^Q2058R^ (residues 2036-2406) and Ig1921 ^K2260R^ (residues 2036-2406) of the human FLNc gene (UniProt code Q14315) were cloned into p3NH-vector (58) which confers resistance to chloramphenicol and kanamycin, and attaches an N-terminal His_6_ tag followed by a human rhinovirus 3C protease cleavage site.

### Protein expression

For protein expression, 6x 500 mL Autoinduction media (no trace metal mix added) supplemented with 100 µg/mL chloramphenicol and 50 µg/mL kanamycin was inoculated with 20 mL overnight culture and grown at 37°C until OD_600_ ∼0.8 shaking at 150 rpm. Then the temperature was reduced to 20°C and after 12 hours the culture was centrifuged (4°C, 5000xg) and the resulting cell pellet was either processed immediately or frozen in liquid nitrogen and stored at −80°C.

### Protein purification

Ig21 and Ig19 cell pellets were resuspended in 100 mL lysis buffer (1x PBS, 20 mM imidazole, pH 7.4) while Ig20-21, Ig19-21, Ig19-21^Q2058R^, Ig19-21^I2160F^, Ig19-21^K2260R^, Ig19-21^P^ ^2298S^, Ig19-21^W2164C^ cell pellets were resuspended in 100 mL lysis buffer (1 X PBS, 20 mM imidazole, 2 M urea, pH 7.4). Ig14, Ig15, Ig14-15, Ig14-15^S1624L^, Ig14-15^G1676R^ cell pellets were resuspended in 100 mL lysis buffer (50 mM MES, 150 mM NaCl, 2 M urea, pH 6.5). All lysis buffers were supplemented with 50 uL DNaseI (10 mg/mL) and sonicated twice for 3 min (50% Amplitude; 1s Pulse On; 1s Pulse Off). The lysate was clarified by centrifugation and subsequently, the supernatant was loaded onto a 5 mL HisTrap FF crude (GE Healthcare) column pre-equilibrated with lysis buffer. Constructs eluted with a step gradient using an elution buffer (1 X PBS, 500 mM imidazole, pH 7.4) for Ig19 and Ig21, while Ig20-21, Ig19-21, Ig19-21^Q2058R^, Ig19-21^I2160F^, Ig19-21^K2260R^, Ig19-21^P^ ^2298S^ were eluted with 1 X PBS, 500 mM imidazole, 2 M urea, pH 7.4. Ig14, Ig15, Ig14-15, Ig14-15^S1624L^, Ig14-15^G1676R^ were eluted with 50 mM MES, 500 mM imidazole, 2 M urea, pH 6.5. The N-terminal His-tag of Ig14, Ig15, Ig14-15, Ig14-15^S1624L^, Ig14-15^G1676R^ Ig19 and Ig21 was cleaved by overnight incubating with human rhinovirus 3C protease with human rhinovirus 3C protease and afterwards loaded again onto a 5 mL HisTrap FF crude column pre-equilibrated with corresponding lysis buffer. These constructs were concentrated after pooling the fractions containing pure protein. Individual fractions of purified constructs were frozen in liquid nitrogen without further concentration and stored at −80°C until further use. Prior to the DSC experiments, the samples were overnight inoculated with human rhinovirus 3C protease and afterwards loaded again onto a 5 mL HisTrap FF crude column pre-equilibrated with corresponding lysis buffer. The successfully cleaved samples were subsequently buffer exchanged using a Superdex S200 5/150 GL or 10/300 increase column (GE Healthcare) connected to an Agilent HPLC 1260 Affinity equipped with a fraction collector. The column was either equilibrated with 50 mM MES 150 mM NaCl pH 6.5 for Ig14, Ig15, Ig14-15, Ig14-15^G1676R^, Ig14-15^S1624L^ or 1x PBS 150 mM NaCl pH 7.4 for Ig19, Ig20-21 Ig21, Ig19-21, Ig19-21^Q2058R^, Ig19-21^I2160F^, Ig19-21^K2260R^, Ig19-21^P^ ^2298S^ and Ig19-21^W2164C^ constructs.

### Crystallization of Ig14-15, Ig14-15^S1624L^ and Ig14-15^G1676R^

Crystallization was performed using SWISSCI MRC three-well crystallization plates (Molecular Dimensions) applying the sitting drop vapour diffusion method. Crystallization plates with commercially available screens were set up using the Mosquito crystallization robot (TTP LabTech). The reservoir was filled with 35 µL of precipitant solution and different ratios of protein to precipitant (150:200 nL, 200:200 nL and 250:200 nL) were applied. Crystallization plates were stored in a Formulatrix RI-1000 imaging device at 22°C. Crystals of Ig14-15 were obtained in SG1™ (ShotGun) screen (Molecular dimensions) in well G1 (0.1 M HEPES pH 7.0, 30 % v/v Jeffamine® ED-2003). Crystals of Ig14-15S^1624L^ were obtained in PACT premier™ screens (Molecular dimensions) in well A2 (0.1 M SPG buffer: succinic acid, sodium phosphate monobasic monohydrate, glycine; pH 5.0, 25 % w/v PEG 1500) and crystals of Ig14-15^G1676R^ in 0.1 M Bis-TRIS pH 6, 26.4 % w/v PEG 3350, 5% glycerol (initial conditions SG1 condition F11).

Crystals of Ig14-15, Ig14-15^S1624L^ were soaked with mother liquor supplemented with 20% 2-methyl-2,4-pentanediol (MPD), collected with cryo-loops and flash-vitrified with liquid nitrogen. For Ig14-15^G1676R^ mother liquor supplemented with 20% of glycerol was used as cryoprotectant.

### Data collection and structure refinement

Datasets of Ig14-15 were collected at beamline i24 of the Diamond Light Source (UK) at 100 K using a Pilatus3 6M. Crystals of Ig14-15^S1624L^ on i04 of the Diamond Light Source at 100 K using a Pilatus 6M-F and crystals of Ig14-15^G1676R^ at beamline MASSIF-1 at ESRF using a Pilatus3 2M (http://dx.doi.org/10.1107/S1399004715011918). The dataset was processed with XDS and symmetry equivalent reflections merged with XDSCONV (59). Intensities were not converted to amplitudes. Initially, we used a conservative high-resolution cutoff 1.7 Å (CC_1/2_ = 67.5; I/σ(I)=1.57) for Ig14-15, 1.92 Å (CC_1/2_ = 86; I/σ(I)=2.1) for Ig14-15^S1624L^ and 1.83 Å (CC_1/2_ = 93.9; I/σ(I)=3.81) for Ig14-15^G1676R^ (60). The phase problem of Ig14-15 was solved by molecular replacement using MORDA from ccp4 online (61,62). For Ig14-15^S1624L^ and Ig14-15^G1676R^ we used phenix.phaser (63) taking Ig14-15 as the search model. The models were further improved by iterative cycles of a manual model building using COOT (64) and maximum likelihood refinement using phenix.refine (65). Phenix.refine converted intensities into amplitudes using the French and Wilson algorithm (66). The final high-resolution cutoff was based on performing paired refinement using PAIREF (67) and PDB_REDO webserver (68).

Final stages of refinement included translation-libration-screw (TLS) parameters, isotropic B-factor model, automated addition of hydrogens and water molecules, optimization of X-ray/ADP weight, and optimization of X-ray/stereochemistry weight for Ig14-15^S1624L^ and Ig14-15^G1676R^. For Ig14-15 an anisotropic B-factor model. The models were validated with MolProbity (69) and the PDB_REDO webserver. The statistics on data-collection and refinement are reported in **<u>Table S2</u>**.

Figures were prepared with PyMOL Molecular Graphics System (Version 2.4.0, Schrödinger, LLC). Atomic coordinates have been deposited in the Protein Data Bank under the accession code Ig14-15 (7OUU), Ig14-15^S1624L^ (7OUV) and Ig14-15^G1676R^ (7P0E).

### DSC analysis

Based on experimental buffer screening (not shown here), MES buffer (50 mM MES, 150 mM NaCl pH 6.5) was selected for Ig14-15 constructs and PBS (1x PBS, 150 mM NaCl pH 7.4) buffer for Ig19-21 constructs. The constructs were investigated in the temperature range of 20 to 110°C. Protein concentration was determined by 280 nm absorbance. The extinction coefficients were calculated from the primary amino acid sequence using Protparam (https://web.expasy.org/protparam).

Both buffers are DSC compatible and did not show any extensive contribution to heat capacity which could distort results. We kept concentrations of wild-types and mutants around 1 mg/ml. Due to ongoing aggregation during concentration, Ig19-21^I2160F^ and Ig19-21^P2298S^ were measured at 0.82 mg/ml and 0.67 mg/ml, respectively, which still yielded interpretable thermograms. Normalized values were used for comparison of different mutations in terms of changes in enthalpy upon unfolding. For data analysis, a “non two state” model was employed to fit all thermograms, besides Ig19, Ig14, Ig15 and Ig21, where a “two state” model was used to fit the experimental data based on agreement between calculated Van’t Hof and observed calorimetric enthalpy. For each experimental run, the baseline approximation under the transition curve was calculated with a spline approximation. Prior to each experimental construct run, we conducted at least three buffer runs to counteract thermal hysteresis of the device and chose the most stable buffer run for later buffer subtraction. All experiments from this study were conducted on a MicroCal PEAQ-DSC from Malvern Panalytical.

## Web-server, data and code availability

AMIVA-F is accessible as a web server based version through http://amiva.msp.univie.ac.at/ which does not require any mandatory dependencies.

A standalone version of AMIVA-F can be found under https://pypi.org/project/AMIVA-F/ or https://github.com/nagym72/AMIVA-F.

The atomic coordinates of all homology models (and AlphaFold2 validation structures) are available upon request or downloadable from PyPi under the project folder of AMIVA-F (https://pypi.org/project/AMIVA-F/). The published package contains a training set, homology and experimentally derived structures and tutorials.

While the web server runs without any dependencies, an in depth tutorial guiding the installation process and the usage of AMIVA-F for the standalone version is included. In order to utilize the standalone AMIVA-F version, a Java virtual machine (JVM) and PyMol are required (instructions on installation for a step by step guidance is specified at https://pypi.org/project/AMIVA-F/).

Java is required in order to access WEKA, the machine learning platform, while Pymol is required to compute some of the input parameters required for WEKA.

Inside the AMIVA-F GUI, an extensive tutorial and additional information for advanced users is available. AMIVA-F is tested on Windows10, as well as in a virtual environment (Anaconda 3, from python=3.6 up to 3.9), MacOSF and Linux (Ubuntu 20.04.2 LTS, Ubuntu 22.04.1) and is designed to be operating system independent.

The crystal structures have been deposited in the Protein Data Bank and are available with these links and will be released upon publication:

https://www.rcsb.org/structure/unreleased/7OUU

https://www.rcsb.org/structure/unreleased/7OUV

https://www.rcsb.org/structure/unreleased/7P0E

## Web resources

*Please provide a URL and title for the website at which the novel computer program described in the manuscript will be made publicly available in a Web Resources section within the manuscript*.

Website at which AMIVA-F is publicly available: amiva.msp.univie.ac.at https://pypi.org/project/AMIVA-F/

## Author contributions

Conceptualization: KD-C, OC; investigation: MN, GM, LS, MMA, LRL; methodology: MN, GM, DP, LRL; visualization: MN, JK, MG; data curation: KD-C, OC; supervision: KD-C, OC, PME, MG, OC, KD-C; validation: KD-C, OC; resources: KD-C, OC, DOF, PC, TBR, ZB, PS, PME, MG; funding acquisition: KD-C, MG, PME; writing (original draft):MN, KD-C, OC; writing (review and editing): all authors.

## Acknowledgements

This work was supported by the Slovenian Research Agency young researcher grant (No. 35337) and research program P1-0140. KDC research was supported by a Marie Curie Initial Training Network: MUZIC (N°238423), Austrian Science Fund (FWF) Projects I525, I1593, P22276, P19060 and W1221, Federal Ministry of Economy, Family and Youth through the initiative “Laura Bassi Centres of Expertise” funding the Centre of Optimized Structural Studies, N°253275, by the Wellcome Trust Collaborative Award (201543/Z/16), Austrian-Slovak Interreg Project B301 StruBioMol, COST action BM1405 - Non-globular proteins - from sequence to structure, function and application in molecular physiopathology (NGP-NET), WWTF (Vienna Science and Technology Fund) Chemical Biology project LS17-008, and by the University of Vienna. JK was supported by the Wellcome Trust Collaborative Award (201543/Z/16) and Austrian-Slovak Interreg Project B301 StruBioMol, LS was supported by the Wellcome Trust Collaborative Award (201543/Z/16). MG holds the British Heart Foundation Chair of Molecular Cardiology.

We thank Jürgen Hoffmann (UNIVIE) aiding in testing and setting up AMIVA-F for MacOS and several Linux partitions.

OC acknowledges support from the Ministero dell’Università e della Ricerca (MUR) and the University of Pavia through the program “Dipartimenti di Eccellenza 2023–2027”. OC also thanks G. Frescobaldi for constant support.

LRL is funded by an MRC UK Clinical Academic Research Partnership (MR/T005181/1).

PC is funded by AVIESAN-ITMO Genetique-Genomique-Bioinformatique [ResDiCard project, Rare diseases call].

The authors would like to thank Diamond Light Source for beamtime (proposal mx20221), and the staff of beamlines I04, and I24 for assistance with crystal testing and data collection.

The access to DSC equipment was kindly provided by the EQ-BOKU VIBT GmbH and the BOKU Core Facility Biomolecular & Cellular Analysis. We thank Jakob Wallner for support.

Furthermore, we thank “Team: Webinterface AMIVA-F Program”, consisting of Stefan Burghuber, Bernd Cala, Antonia Schwarz, Benjamin Wittmann during their final year at the “Kolleg für Informatik” at HTL Spengergasse, Vienna.

## References

1. Disease, G. B. D., Injury, I., and Prevalence, C. (2018) Global, regional, and national incidence, prevalence, and years lived with disability for 354 diseases and injuries for 195 countries and territories, 1990-2017: a systematic analysis for the Global Burden of Disease Study 2017. Lancet 392, 1789–1858; 10.1016/S0140-6736(18)32279-7

2. Elliott, P., Andersson, B., Arbustini, E., Bilinska, Z., Cecchi, F., Charron, P., Dubourg, O., Kuhl, U., Maisch, B., McKenna, W. J., Monserrat, L., Pankuweit, S., Rapezzi, C., Seferovic, P., Tavazzi, L., and Keren, A. (2008) Classification of the cardiomyopathies: a position statement from the European Society Of Cardiology Working Group on Myocardial and Pericardial Diseases. Eur Heart J 29, 270–276; 10.1093/eurheartj/ehm342

3. Agarwal, R., Paulo, J. A., Toepfer, C. N., Ewoldt, J. K., Sundaram, S., Chopra, A., Zhang, Q., Gorham, J., Depalma, S. R., Chen, C. S., Gygi, S. P., Seidman, C. E., and Seidman, J. G. (2021) Filamin C Cardiomyopathy Variants Cause Protein and Lysosome Accumulation. Circulation Research 129, 751–766; 10.1161/circresaha.120.317076

4. Verdonschot, J. A. J., Vanhoutte, E. K., Claes, G. R. F., Helderman-van den Enden, A., Hoeijmakers, J. G. J., Hellebrekers, D., de Haan, A., Christiaans, I., Lekanne Deprez, R. H., Boen, H. M., van Craenenbroeck, E. M., Loeys, B. L., Hoedemaekers, Y. M., Marcelis, C., Kempers, M., Brusse, E., van Waning, J. I., Baas, A. F., Dooijes, D., Asselbergs, F. W., Barge-Schaapveld, D., Koopman, P., van den Wijngaard, A., Heymans, S. R. B., Krapels, I. P. C., and Brunner, H. G. (2020) A mutation update for the FLNC gene in myopathies and cardiomyopathies. Hum Mutat 41, 1091–1111; 10.1002/humu.24004

5. Valdes-Mas, R., Gutierrez-Fernandez, A., Gomez, J., Coto, E., Astudillo, A., Puente, D. A., Reguero, J. R., Alvarez, V., Moris, C., Leon, D., Martin, M., Puente, X. S., and Lopez-Otin, C. (2014) Mutations in filamin C cause a new form of familial hypertrophic cardiomyopathy. Nat Commun 5, 5326; 10.1038/ncomms6326

6. Ortiz-Genga, M. F., Cuenca, S., Dal Ferro, M., Zorio, E., Salgado-Aranda, R., Climent, V., Padrón-Barthe, L., Duro-Aguado, I., Jiménez-Jáimez, J., Hidalgo-Olivares, V. M., García-Campo, E., Lanzillo, C., Suárez-Mier, M. P., Yonath, H., Marcos-Alonso, S., Ochoa, J. P., Santomé, J. L., García-Giustiniani, D., Rodríguez-Garrido, J. L., Domínguez, F., Merlo, M., Palomino, J., Peña, M. L., Trujillo, J. P., Martín-Vila, A., Stolfo, D., Molina, P., Lara-Pezzi, E., Calvo-Iglesias, F. E., Nof, E., Calò, L., Barriales-Villa, R., Gimeno-Blanes, J. R., Arad, M., García-Pavía, P., and Monserrat, L. (2016) Truncating FLNC Mutations Are Associated With High-Risk Dilated and Arrhythmogenic Cardiomyopathies. Journal of the American College of Cardiology 68, 2440–2451; 10.1016/j.jacc.2016.09.927

7. Ader, F., De Groote, P., Réant, P., Rooryck-Thambo, C., Dupin-Deguine, D., Rambaud, C., Khraiche, D., Perret, C., Pruny, J. F., Mathieu-Dramard, M., Gérard, M., Troadec, Y., Gouya, L., Jeunemaitre, X., Van Maldergem, L., Hagège, A., Villard, E., Charron, P., and Richard, P. (2019) FLNC pathogenic variants in patients with cardiomyopathies: Prevalence and genotype-phenotype correlations. Clinical Genetics 96, 317–329; 10.1111/cge.13594

8. Eden, M., and Frey, N. (2021) Cardiac Filaminopathies: Illuminating the Divergent Role of Filamin C Mutations in Human Cardiomyopathy. Journal of Clinical Medicine 10, 577; 10.3390/jcm10040577

9. Wadmore, K., Azad, A. J., and Gehmlich, K. (2021) The Role of Z-disc Proteins in Myopathy and Cardiomyopathy. International Journal of Molecular Sciences 22, 3058; 10.3390/ijms22063058

10. Thompson, T. G., Chan, Y. M., Hack, A. A., Brosius, M., Rajala, M., Lidov, H. G., McNally, E. M., Watkins, S., and Kunkel, L. M. (2000) Filamin 2 (FLN2): A muscle-specific sarcoglycan interacting protein. J Cell Biol 148, 115–126; 10.1083/jcb.148.1.115

11. Mao, Z., and Nakamura, F. (2020) Structure and Function of Filamin C in the Muscle Z-Disc. Int J Mol Sci 2110.3390/ijms21082696

12. Fujita, M., Mitsuhashi, H., Isogai, S., Nakata, T., Kawakami, A., Nonaka, I., Noguchi, S., Hayashi, Y. K., Nishino, I., and Kudo, A. (2012) Filamin C plays an essential role in the maintenance of the structural integrity of cardiac and skeletal muscles, revealed by the medaka mutant zacro. Developmental Biology 361, 79–89; 10.1016/j.ydbio.2011.10.008

13. Leber, Y., Ruparelia, A. A., Kirfel, G., van der Ven, P. F., Hoffmann, B., Merkel, R., Bryson-Richardson, R. J., and Furst, D. O. (2016) Filamin C is a highly dynamic protein associated with fast repair of myofibrillar microdamage. Hum Mol Genet 25, 2776–2788; 10.1093/hmg/ddw135

14. Molt, S., Buhrdel, J. B., Yakovlev, S., Schein, P., Orfanos, Z., Kirfel, G., Winter, L., Wiche, G., van der Ven, P. F., Rottbauer, W., Just, S., Belkin, A. M., and Furst, D. O. (2014) Aciculin interacts with filamin C and Xin and is essential for myofibril assembly, remodeling and maintenance. J Cell Sci 127, 3578–3592; 10.1242/jcs.152157

15. van der Flier, A., Kuikman, I., Kramer, D., Geerts, D., Kreft, M., Takafuta, T., Shapiro, S. S., and Sonnenberg, A. (2002) Different splice variants of filamin-B affect myogenesis, subcellular distribution, and determine binding to integrin [beta] subunits. J Cell Biol 156, 361–376; 10.1083/jcb.200103037

16. Lad, Y., Kiema, T., Jiang, P., Pentikainen, O. T., Coles, C. H., Campbell, I. D., Calderwood, D. A., and Ylanne, J. (2007) Structure of three tandem filamin domains reveals auto-inhibition of ligand binding. EMBO J 26, 3993–4004; 10.1038/sj.emboj.7601827

17. Sethi, R., and Ylänne, J. (2014) Small-Angle X-Ray Scattering Reveals Compact Domain-Domain Interactions in the N-Terminal Region of Filamin C. PLoS ONE 9, e107457; 10.1371/journal.pone.0107457

18. Seppälä, J., Bernardi, R. C., Haataja, T. J. K., Hellman, M., Pentikäinen, O. T., Schulten, K., Permi, P., Ylänne, J., and Pentikäinen, U. (2017) Skeletal Dysplasia Mutations Effect on Human Filamins’ Structure and Mechanosensing. Scientific Reports 710.1038/s41598-017-04441-x

19. Tossavainen, H., Koskela, O., Jiang, P., Ylänne, J., Campbell, I. D., Kilpeläinen, I., and Permi, P. (2012) Model of a Six Immunoglobulin-Like Domain Fragment of Filamin A (16–21) Built Using Residual Dipolar Couplings. Journal of the American Chemical Society 134, 6660–6672; 10.1021/ja2114882

20. Ruskamo, S., Gilbert, R., Hofmann, G., Jiang, P., Campbell, I. D., Ylanne, J., and Pentikainen, U. (2012) The C-terminal rod 2 fragment of filamin A forms a compact structure that can be extended. Biochem J 446, 261–269; 10.1042/BJ20120361

21. Ehrlicher, A. J., Nakamura, F., Hartwig, J. H., Weitz, D. A., and Stossel, T. P. (2011) Mechanical strain in actin networks regulates FilGAP and integrin binding to filamin A. Nature 478, 260–263; 10.1038/nature10430

22. Rognoni, L., Stigler, J., Pelz, B., Ylanne, J., and Rief, M. (2012) Dynamic force sensing of filamin revealed in single-molecule experiments. Proc Natl Acad Sci U S A 109, 19679–19684; 10.1073/pnas.1211274109

23. Rognoni, L., Most, T., Zoldak, G., and Rief, M. (2014) Force-dependent isomerization kinetics of a highly conserved proline switch modulates the mechanosensing region of filamin. Proc Natl Acad Sci U S A 111, 5568–5573; 10.1073/pnas.1319448111

24. Brodehl, A., Ferrier, R. A., Hamilton, S. J., Greenway, S. C., Brundler, M.-A., Yu, W., Gibson, W. T., McKinnon, M. L., McGillivray, B., Alvarez, N., Giuffre, M., Schwartzentruber, J., Consortium, F. C., and Gerull, B. (2016) Mutations in FLNC are Associated with Familial Restrictive Cardiomyopathy. Human Mutation 37, 269–279; https://doi.org/10.1002/humu.22942

25. Reimann, L., Wiese, H., Leber, Y., Schwable, A. N., Fricke, A. L., Rohland, A., Knapp, B., Peikert, C. D., Drepper, F., van der Ven, P. F., Radziwill, G., Furst, D. O., and Warscheid, B. (2017) Myofibrillar Z-discs Are a Protein Phosphorylation Hot Spot with Protein Kinase C (PKCalpha) Modulating Protein Dynamics. Mol Cell Proteomics 16, 346–367; 10.1074/mcp.M116.065425

26. Reimann, L., Schwable, A. N., Fricke, A. L., Muhlhauser, W. W. D., Leber, Y., Lohanadan, K., Puchinger, M. G., Schauble, S., Faessler, E., Wiese, H., Reichenbach, C., Knapp, B., Peikert, C. D., Drepper, F., Hahn, U., Kreutz, C., van der Ven, P. F. M., Radziwill, G., Djinovic-Carugo, K., Furst, D. O., and Warscheid, B. (2020) Phosphoproteomics identifies dual-site phosphorylation in an extended basophilic motif regulating FILIP1-mediated degradation of filamin-C. Commun Biol 3, 253; 10.1038/s42003-020-0982-5

27. Pires, D. E. V., Ascher, D. B., and Blundell, T. L. (2014) DUET: a server for predicting effects of mutations on protein stability using an integrated computational approach. Nucleic Acids Research 42, W314–W319; 10.1093/nar/gku411

28. Gómez, J., Lorca, R., Reguero, J. R., Morís, C., Martín, M., Tranche, S., Alonso, B., Iglesias, S., Alvarez, V., Díaz-Molina, B., Avanzas, P., and Coto, E. (2017) Screening of the Filamin C Gene in a Large Cohort of Hypertrophic Cardiomyopathy Patients. Circulation: Cardiovascular Genetics 10, e001584; 10.1161/circgenetics.116.001584

29. Ruskamo, S., Gilbert, R., Hofmann, G., Jiang, P., Iain, Ylänne, J., and Pentikäinen, U. (2012) The C-terminal rod 2 fragment of filamin A forms a compact structure that can be extended. Biochemical Journal 446, 261–269<otherinfo>; 10.1042/bj20120361</otherinfo>

30. Roy, A., Kucukural, A., and Zhang, Y. (2010) I-TASSER: a unified platform for automated protein structure and function prediction. Nature Protocols 5, 725–738; 10.1038/nprot.2010.5

31. Yang, J., and Zhang, Y. (2015) I-TASSER server: new development for protein structure and function predictions. Nucleic Acids Research 43, W174–W181; 10.1093/nar/gkv342

32. Ashkenazy, H., Abadi, S., Martz, E., Chay, O., Mayrose, I., Pupko, T., and Ben-Tal, N. (2016) ConSurf 2016: an improved methodology to estimate and visualize evolutionary conservation in macromolecules. Nucleic Acids Research 44, W344–W350; 10.1093/nar/gkw408

33. Karczewski, K. J., Francioli, L. C., Tiao, G., Cummings, B. B., Alföldi, J., Wang, Q., Collins, R. L., Laricchia, K. M., Ganna, A., Birnbaum, D. P., Gauthier, L. D., Brand, H., Solomonson, M., Watts, N. A., Rhodes, D., Singer-Berk, M., England, E. M., Seaby, E. G., Kosmicki, J. A., Walters, R. K., Tashman, K., Farjoun, Y., Banks, E., Poterba, T., Wang, A., Seed, C., Whiffin, N., Chong, J. X., Samocha, K. E., Pierce-Hoffman, E., Zappala, Z., O’Donnell-Luria, A. H., Minikel, E. V., Weisburd, B., Lek, M., Ware, J. S., Vittal, C., Armean, I. M., Bergelson, L., Cibulskis, K., Connolly, K. M., Covarrubias, M., Donnelly, S., Ferriera, S., Gabriel, S., Gentry, J., Gupta, N., Jeandet, T., Kaplan, D., Llanwarne, C., Munshi, R., Novod, S., Petrillo, N., Roazen, D., Ruano-Rubio, V., Saltzman, A., Schleicher, M., Soto, J., Tibbetts, K., Tolonen, C., Wade, G., Talkowski, M. E., Neale, B. M., Daly, M. J., and Macarthur, D. G. (2020) The mutational constraint spectrum quantified from variation in 141,456 humans. Nature 581, 434–443; 10.1038/s41586-020-2308-7

34. van der Ven, P. F., Ehler, E., Vakeel, P., Eulitz, S., Schenk, J. A., Milting, H., Micheel, B., and Furst, D. O. (2006) Unusual splicing events result in distinct Xin isoforms that associate differentially with filamin c and Mena/VASP. Exp Cell Res 312, 2154–2167; 10.1016/j.yexcr.2006.03.015

35. Worth, C. L., Preissner, R., and Blundell, T. L. (2011) SDM--a server for predicting effects of mutations on protein stability and malfunction. Nucleic Acids Research 39, W215–W222; 10.1093/nar/gkr363

36. Kyte, J., and Doolittle, R. F. (1982) A simple method for displaying the hydropathic character of a protein. J Mol Biol 157, 105–132; 10.1016/0022-2836(82)90515-0

37. Chennamsetty, N., Voynov, V., Kayser, V., Helk, B., and Trout, B. L. (2009) Design of therapeutic proteins with enhanced stability. Proceedings of the National Academy of Sciences 106, 11937–11942; 10.1073/pnas.0904191106

38. Witten, I. H., Frank, E., Hall, M. A., and Pal, C. J. (2017) Data Mining: Practical Machine Learning Tools and Techniques, 4th Edition. Data Mining: Practical Machine Learning Tools and Techniques, 4th Edition, Cp1-621;

39. Matthews, B. W. (1975) Comparison of the predicted and observed secondary structure of T4 phage lysozyme. Biochim Biophys Acta 405, 442–451; 10.1016/0005-2795(75)90109-9

40. Fokkema, I. F., Taschner, P. E., Schaafsma, G. C., Celli, J., Laros, J. F., and den Dunnen, J. T. (2011) LOVD v.2.0: the next generation in gene variant databases. Hum Mutat 32, 557-563; 10.1002/humu.21438

41. Richards, S., Aziz, N., Bale, S., Bick, D., Das, S., Gastier-Foster, J., Grody, W. W., Hegde, M., Lyon, E., Spector, E., Voelkerding, K., Rehm, H. L., and Committee, A. L. Q. A. (2015) Standards and guidelines for the interpretation of sequence variants: a joint consensus recommendation of the American College of Medical Genetics and Genomics and the Association for Molecular Pathology. Genet Med 17, 405–424; 10.1038/gim.2015.30

42. Lek, M., Karczewski, K. J., Minikel, E. V., Samocha, K. E., Banks, E., Fennell, T., O’Donnell-Luria, A. H., Ware, J. S., Hill, A. J., Cummings, B. B., Tukiainen, T., Birnbaum, D. P., Kosmicki, J. A., Duncan, L. E., Estrada, K., Zhao, F., Zou, J., Pierce-Hoffman, E., Berghout, J., Cooper, D. N., Deflaux, N., Depristo, M., Do, R., Flannick, J., Fromer, M., Gauthier, L., Goldstein, J., Gupta, N., Howrigan, D., Kiezun, A., Kurki, M. I., Moonshine, A. L., Natarajan, P., Orozco, L., Peloso, G. M., Poplin, R., Rivas, M. A., Ruano-Rubio, V., Rose, S. A., Ruderfer, D. M., Shakir, K., Stenson, P. D., Stevens, C., Thomas, B. P., Tiao, G., Tusie-Luna, M. T., Weisburd, B., Won, H.-H., Yu, D., Altshuler, D. M., Ardissino, D., Boehnke, M., Danesh, J., Donnelly, S., Elosua, R., Florez, J. C., Gabriel, S. B., Getz, G., Glatt, S. J., Hultman, C. M., Kathiresan, S., Laakso, M., McCarroll, S., McCarthy, M. I., McGovern, D., McPherson, R., Neale, B. M., Palotie, A., Purcell, S. M., Saleheen, D., Scharf, J. M., Sklar, P., Sullivan, P. F., Tuomilehto, J., Tsuang, M. T., Watkins, H. C., Wilson, J. G., Daly, M. J., and Macarthur, D.G. (2016) Analysis of protein-coding genetic variation in 60,706 humans. Nature 536, 285–291; 10.1038/nature19057

43. Selcen, D., Ohno, K., and Engel, A. G. (2004) Myofibrillar myopathy: clinical, morphological and genetic studies in 63 patients. Brain 127, 439–451; 10.1093/brain/awh052

44. Ader, F., De Groote, P., Réant, P., Rooryck-Thambo, C., Dupin-Deguine, D., Rambaud, C., Khraiche, D., Perret, C., Pruny, J. F., Mathieu-Dramard, M., Gérard, M., Troadec, Y., Gouya, L., Jeunemaitre, X., Van Maldergem, L., Hagège, A., Villard, E., Charron, P., and Richard, P. (2019) FLNC pathogenic variants in patients with cardiomyopathies: Prevalence and genotype-phenotype correlations. Clinical Genetics 96, 317–329; https://doi.org/10.1111/cge.13594

45. Akhtar, M. M., Lorenzini, M., Pavlou, M., Ochoa, J. P., O’Mahony, C., Restrepo-Cordoba, M. A., Segura-Rodriguez, D., Bermudez-Jimenez, F., Molina, P., Cuenca, S., Ader, F., Larranaga-Moreira, J. M., Sabater-Molina, M., Garcia-Alvarez, M. I., Arantzamendi, L. G., Truszkowska, G., Ortiz-Genga, M., Ruiz, I. S., Nielson, S. K., Rasmussen, T. B., Robles Mezcua, A., Alvarez-Rubio, J., Eiskjaer, H., Gautel, M., Garcia-Pinilla, J. M., Ripoll-Vera, T., Mogensen, J., Limeres Freire, J., Rodriguez-Palomares, J. F., Pena-Pena, M. L., Rangel-Sousa, D., Palomino-Doza, J., Arana Achaga, X., Bilinska, Z., Zamarreno Golvano, E., Climent, V., Penalver, M. N., Barriales-Villa, R., Charron, P., Yotti, R., Zorio, E., Jimenez-Jaimez, J., Garcia-Pavia, P., Elliott, P. M., and European Genetic Cardiomyopathies Initiative, I. (2021) Association of Left Ventricular Systolic Dysfunction Among Carriers of Truncating Variants in Filamin C With Frequent Ventricular Arrhythmia and End-stage Heart Failure. JAMA Cardiol 10.1001/jamacardio.2021.1106

46. Song, S., Shi, A., Lian, H., Hu, S., and Nie, Y. (2022) Filamin C in cardiomyopathy: from physiological roles to DNA variants. Heart Fail Rev 27, 1373–1385; 10.1007/s10741-021-10172-z

47. Stewart, D. E., Sarkar, A., and Wampler, J. E. (1990) Occurrence and role of cis peptide bonds in protein structures. J Mol Biol 214, 253–260; 10.1016/0022-2836(90)90159-J

48. Adzhubei, I. A., Schmidt, S., Peshkin, L., Ramensky, V. E., Gerasimova, A., Bork, P., Kondrashov, A. S., and Sunyaev, S.R. (2010) A method and server for predicting damaging missense mutations. Nature Methods 7, 248–249; 10.1038/nmeth0410-248

49. Ng, P. C. (2003) SIFT: predicting amino acid changes that affect protein function. Nucleic Acids Research 31, 3812–3814; 10.1093/nar/gkg509

50. Choi, Y., Sims, G. E., Murphy, S., Miller, J. R., and Chan, A. P. (2012) Predicting the Functional Effect of Amino Acid Substitutions and Indels. PLoS ONE 7, e46688; 10.1371/journal.pone.0046688

51. Lange, S., Gehmlich, K., Lun, A. S., Blondelle, J., Hooper, C., Dalton, N. D., Alvarez, E. A., Zhang, X., Bang, M. L., Abassi, Y. A., Dos Remedios, C. G., Peterson, K. L., Chen, J., and Ehler, E. (2016) MLP and CARP are linked to chronic PKCalpha signalling in dilated cardiomyopathy. Nat Commun 7, 12120; 10.1038/ncomms12120

52. Fürst, D. O., Osborn, M., Nave, R., and Weber, K. (1988) The organization of titin filaments in the half-sarcomere revealed by monoclonal antibodies in immunoelectron microscopy: a map of ten nonrepetitive epitopes starting at the Z line extends close to the M line. Journal of Cell Biology 106, 1563–1572; 10.1083/jcb.106.5.1563

53. Bernstein, F. C., Koetzle, T. F., Williams, G. J., Meyer, E. F., Jr., Brice, M. D., Rodgers, J. R., Kennard, O., Shimanouchi, T., and Tasumi, M. (1977) The Protein Data Bank: a computer-based archival file for macromolecular structures. J Mol Biol 112, 535–542; 10.1016/s0022-2836(77)80200-3

54. Berman, H. M., Westbrook, J., Feng, Z., Gilliland, G., Bhat, T. N., Weissig, H., Shindyalov, I. N., and Bourne, P. E. (2000) The Protein Data Bank. Nucleic Acids Res 28, 235–242; 10.1093/nar/28.1.235

55. Burley, S. K., Berman, H. M., Bhikadiya, C., Bi, C., Chen, L., Di Costanzo, L., Christie, C., Dalenberg, K., Duarte, J. M., Dutta, S., Feng, Z., Ghosh, S., Goodsell, D. S., Green, R. K., Guranovic, V., Guzenko, D., Hudson, B. P., Kalro, T., Liang, Y., Lowe, R., Namkoong, H., Peisach, E., Periskova, I., Prlic, A., Randle, C., Rose, A., Rose, P., Sala, R., Sekharan, M., Shao, C., Tan, L., Tao, Y. P., Valasatava, Y., Voigt, M., Westbrook, J., Woo, J., Yang, H., Young, J., Zhuravleva, M., and Zardecki, C. (2019) RCSB Protein Data Bank: biological macromolecular structures enabling research and education in fundamental biology, biomedicine, biotechnology and energy. Nucleic Acids Res 47, D464–D474; 10.1093/nar/gky1004

56. Sali, A., and Blundell, T. L. (1993) Comparative protein modelling by satisfaction of spatial restraints. J Mol Biol 234, 779–815; 10.1006/jmbi.1993.1626

57. Jumper, J., Evans, R., Pritzel, A., Green, T., Figurnov, M., Ronneberger, O., Tunyasuvunakool, K., Bates, R., Zidek, A., Potapenko, A., Bridgland, A., Meyer, C., Kohl, S. A. A., Ballard, A. J., Cowie, A., Romera-Paredes, B., Nikolov, S., Jain, R., Adler, J., Back, T., Petersen, S., Reiman, D., Clancy, E., Zielinski, M., Steinegger, M., Pacholska, M., Berghammer, T., Bodenstein, S., Silver, D., Vinyals, O., Senior, A. W., Kavukcuoglu, K., Kohli, P., and Hassabis, D. (2021) Highly accurate protein structure prediction with AlphaFold. Nature 596, 583–589; 10.1038/s41586-021-03819-2

58. Mlynek, G., Lehner, A., Neuhold, J., Leeb, S., Kostan, J., Charnagalov, A., Stolt-Bergner, P., Djinović-Carugo, K., and Pinotsis, N. (2014) The Center for Optimized Structural Studies (COSS) platform for automation in cloning, expression, and purification of single proteins and protein-protein complexes. Amino Acids 46, 1565–1582; 10.1007/s00726-014-1699-x

59. Kabsch, W. (2010) XDS. Acta Crystallographica Section D Biological Crystallography 66, 125–132; 10.1107/s0907444909047337

60. Karplus, P. A., and Diederichs, K. (2012) Linking Crystallographic Model and Data Quality. Science 336, 1030–1033; 10.1126/science.1218231

61. Vagin, A., and Lebedev, A. (2015) MoRDa, an automatic molecular replacement pipeline. Acta Crystallographica Section A Foundations and Advances 71, s19–s19; 10.1107/s2053273315099672

62. Winn, M. D., Ballard, C. C., Cowtan, K. D., Dodson, E. J., Emsley, P., Evans, P. R., Keegan, R. M., Krissinel, E. B., Leslie, A. G. W., McCoy, A., McNicholas, S. J., Murshudov, G. N., Pannu, N. S., Potterton, E. A., Powell, H. R., Read, R. J., Vagin, A., and Wilson, K. S. (2011) Overview of theCCP4 suite and current developments. Acta Crystallographica Section D Biological Crystallography 67, 235–242; 10.1107/s0907444910045749

63. McCoy, A. J., Grosse-Kunstleve, R. W., Adams, P. D., Winn, M. D., Storoni, L. C., and Read, R. J. (2007) Phasercrystallographic software. Journal of Applied Crystallography 40, 658–674; 10.1107/s0021889807021206

64. Emsley, P., Lohkamp, B., Scott, W. G., and Cowtan, K. (2010) Features and development ofCoot. Acta Crystallographica Section D Biological Crystallography 66, 486–501; 10.1107/s0907444910007493

65. Adams, P. D., Afonine, P. V., Bunkóczi, G., Chen, V. B., Davis, I. W., Echols, N., Headd, J. J., Hung, L.-W., Kapral, G. J., Grosse-Kunstleve, R. W., McCoy, A. J., Moriarty, N. W., Oeffner, R., Read, R. J., Richardson, D. C., Richardson, J. S., Terwilliger, T. C., and Zwart, P. H. (2010) PHENIX: a comprehensive Python-based system for macromolecular structure solution. Acta Crystallographica Section D Biological Crystallography 66, 213–221; 10.1107/s0907444909052925

66. French, S., and Wilson, K. (1978) Treatment of Negative Intensity Observations. Acta Crystallogr A 34, 517–525; Doi 10.1107/S0567739478001114

67. Malý, M., Diederichs, K., Dohnálek, J., and Kolenko, P. (2020) Paired refinement under the control of PAIREF. IUCrJ 7, 681–692; 10.1107/s2052252520005916

68. Joosten, R. P., Salzemann, J., Bloch, V., Stockinger, H., Berglund, A.-C., Blanchet, C., Bongcam-Rudloff, E., Combet, C., Da Costa, A. L., Deleage, G., Diarena, M., Fabbretti, R., Fettahi, G., Flegel, V., Gisel, A., Kasam, V., Kervinen, T., Korpelainen, E., Mattila, K., Pagni, M., Reichstadt, M., Breton, V., Tickle, I. J., and Vriend, G. (2009) PDB_REDO: automated re-refinement of X-ray structure models in the PDB. Journal of Applied Crystallography 42, 376–384; 10.1107/s0021889809008784

69. Davis, I. W., Murray, L. W., Richardson, J. S., and Richardson, D. C. (2004) MOLPROBITY: structure validation and all-atom contact analysis for nucleic acids and their complexes. Nucleic Acids Res 32, W615–619; 10.1093/nar/gkh398

