## Supplementary information: Text, Figures and Tables for "Unlocking Predictive Power: A Machine Learning Tool Derived from In-Depth Analysis to Forecast the Impact of Missense Variants in Human Filamin C"

**Short title**: A bioinformatics tool for analysis of missense variants in filamin C: AMIVA-F

Michael Nagy[1,#], Georg Mlynek[1,2], Julius Kostan[1], Luke Smith[3], Dominic Pühringer[1], Philippe Charron[4,5], Torsten Bloch Rasmussen[6], Zofia Bilinska[7], Mohammed Majid Akhtar[8,9], Petros Syrris[8], Luis R Lopes[8,9], Perry M Elliott[8,9], Mathias Gautel[3], Oliviero Carugo[1,10,*], Kristina Djinović-Carugo[1,11,12]*

[1] Department of Structural and Computational Biology, Max Perutz Labs, University of Vienna, Vienna, Austria
[2] BOKU Core Facility Biomolecular & Cellular Analysis, BOKU – University of Natural Resources and Life Sciences, Vienna, Austria
[3] King's College London BHF Centre for Research Excellence, Randall Centre for Cell and Molecular Biophysics, London SE1 1UL, UK
[4] APHP, Centre de référence pour les maladies cardiaques héréditaires, Département de Génétique, Hôpital Pitié-Salpêtrière, Paris, France and Sorbonne Université, INSERM, UMRS 1166 and ICAN Institute for Cardiometabolism and Nutrition, Paris, France
[5] Sorbonne Université, INSERM, UMRS 1166 and ICAN Institute for Cardiometabolism and Nutrition, Paris, France
[6] Department of Cardiology, Aarhus University Hospital, Aarhus, Denmark
[7] Unit for Screening Studies in Inherited Cardiovascular Diseases, National Institute of Cardiology, Warsaw, Poland
[8] Center for Heart Muscle Disease, Institute of Cardiovascular Science, University College London, London, UK
[9] Barts Heart Centre, St Bartholomew's Hospital, Barts Health NHS Trust, London, UK
[10] Department of Chemistry, University of Pavia, Pavia, Italy (ORCID 0000-0002-2924-9016)
[11] Department of Biochemistry, Faculty of Chemistry and Chemical Technology, University of Ljubljana, Ljubljana, Slovenia
[12] European Molecular Biology Laboratory (EMBL), Grenoble, France (ORCID 0000-0003-0252-2972)

[#]Current address: Protein Dynamics and Cancer Lab, Department of Oncology-Pathology, Karolinska Institute, Solna, Sweden

### Supplementary Information

#### Pathogenicity propensity upon mutation

The 108 pathogenic variants of human filamin C imply that, on average, there are 4.3 deleterious mutations per domain (one ABD domain and 24 Ig domains). However, the mutations are not equally distributed (**Figure 5).** Some domains are preserved: no pathogenic variants imply mutations in Ig3, Ig9, Ig11 and Ig13 – interestingly, these are odd numbers and might indicate a feeble evolutionary signal. Other domains, on the contrary, are heavily affected, especially in the N-terminal moiety (nine variants imply mutations in the ABD and seven in the Ig1 domain) and in the C-terminal rod-2 region (Ig16-Ig21), where 6.8 deleterious mutations per domain are observed. The rod-2 mutational hotspot is the Ig20 domain, which hosts the mutations implied by 18 pathogenic variants.

Given that the 108 pathogenic variants of human FlnC are associated with a mutation from amino acid R1 to amino acid R2 it is possible to compute the propensity of each residue type to be R1. For example, the propensity of alanine to be R1 (p_Ala_R1) is defined as:

p_Ala_R1 = (n_Ala_R1/108)/(n_Ala/n_res) (equation 1)

where n_Ala_R1 is the number of times alanine is observed in the 108 R1 positions, n_Ala is the number of alanine residues in human FlnC, and n_res = 2725 is the total number of residues in human FlnC (Uniprot Q14315-1). The numerator of equation 1 is the probability to observe an alanine in position R1 amongst the pathogenic variants and the denominator of equation 1 is the probability to observe an alanine in human FlnC. The propensity values are equal to one when the numerator and the denominator are equal, which means that the two probabilities are identical. On the contrary, the propensity is larger than one (or lower than one) if the amino acid tends to produce pathogenic variants upon its mutation (or it does not tend to produce pathogenic variants upon its mutation).

For obvious reasons it is not possible to define the propensity of each residue type to be R2, since the denominator of equation 1 is undetermined in that case.

The 20 propensity values reported in **Supplementary Table S9** are quite interesting. Two amino acids, methionine and arginine, have high propensity to be R1, which means that their mutations are often associated with disease onset.

The high propensity of methionine is likely related to two facts. This residue is usually buried into the protein core and it is unique, with its long aliphatic side-chain, amongst apolar residues. Interestingly, the sulfur atom of the methionine side-chain may be involved in stabilizing chalcogen bonds [1]. Methionine mutations are therefore very likely deleterious for protein stability and, as a consequence, for human FlnC function.

On the contrary, the high propensity to be R1 of arginine is surprising, since this amino acid is usually exposed to the solvent and surface residues of globular proteins are known to be often mutable without major protein destabilization. Moreover, the high propensity of arginine does not compare with that of lysine, which is positively charged like arginine and which has the lowest propensity to be R1 (**Supplementary Table S9**).

Obviously, the sample of pathogenic variants is quite small and it would be temerarious to pretend definitive conclusions from these observations. It is nevertheless possible to hypothesize that arginine methylation, which is involved in numerous and diversified biological processes [2], plays a role in human filamin C function.

Interestingly, in about one third of the mutations from Arg to R2, R2 is a cysteine. This means that replacements of arginines by cysteines have a considerable probability to be pathogenic. This might depend, at least in part, from the fact that solvent accessible cysteines can be easily oxidized and degraded, especially under stress.

#### Disorder/Order analysis in the insertion region of Ig20

Conformational disorder predictions generated with GeneSilico [3] suggest that this insertion is partially disordered (residues 2171-2210) and partially ordered (residues 2210-2243) (**Supplementary S5A**). Similarly, hydrophobic cluster analysis (HCA) [4] predicts for this region islands of hydrophobic clusters after residue 2210 (**Supplementary S5B**). The N-terminal part of IR is rich in Arg, Gly, and Glu residues. High frequency and alternating positive and negative charges are typically observed in intrinsically disordered regions prone to liquid-liquid phase separation and formation of biomolecular condensates [5].

### Supplementary Figures

| **Supplementary Figure S1** Cartoon depiction of FlnC domain structures listed in **Supplementary Table S5 (Structures All)** with associated topologies, highlighting the two planes of ß-sheets in different shades of blue. Ligplot representation of the interface between domains Ig14 and Ig15 observed in the crystal structures of the wild type construct and of the Ig14-15G1676R construct. |
| --- |
| 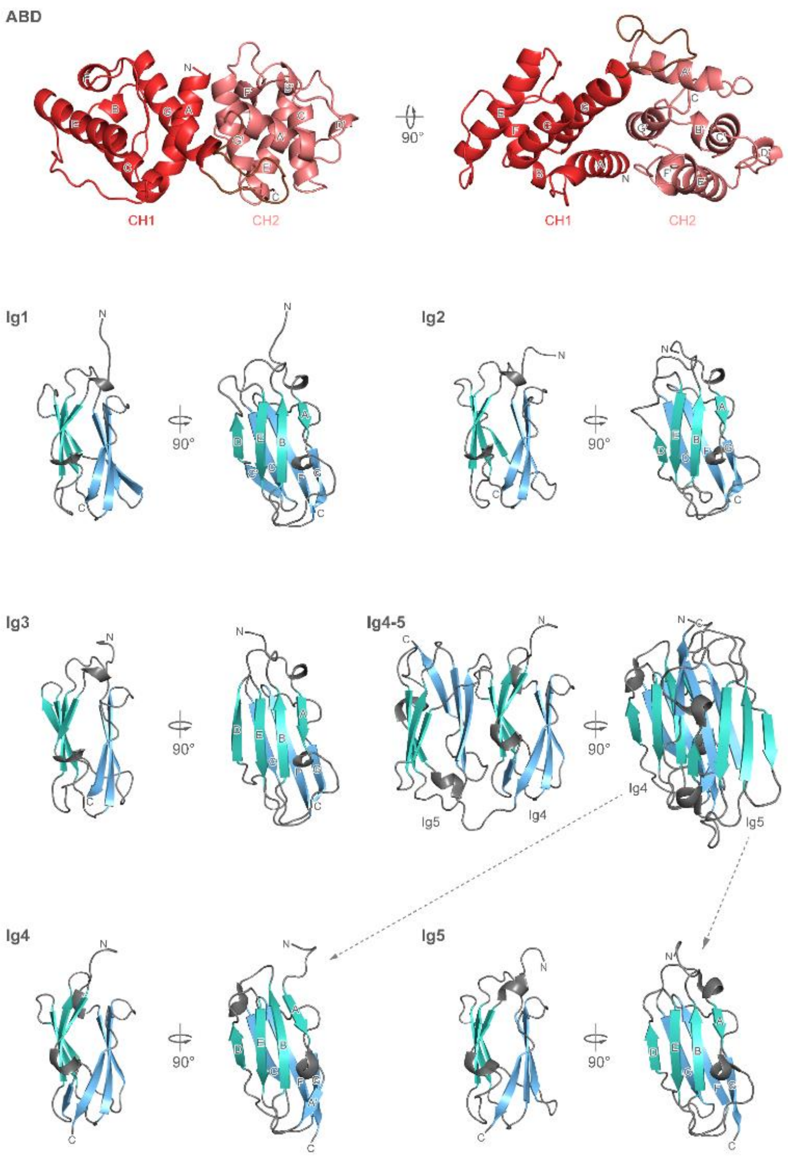 |

**Figure S1** (continue)

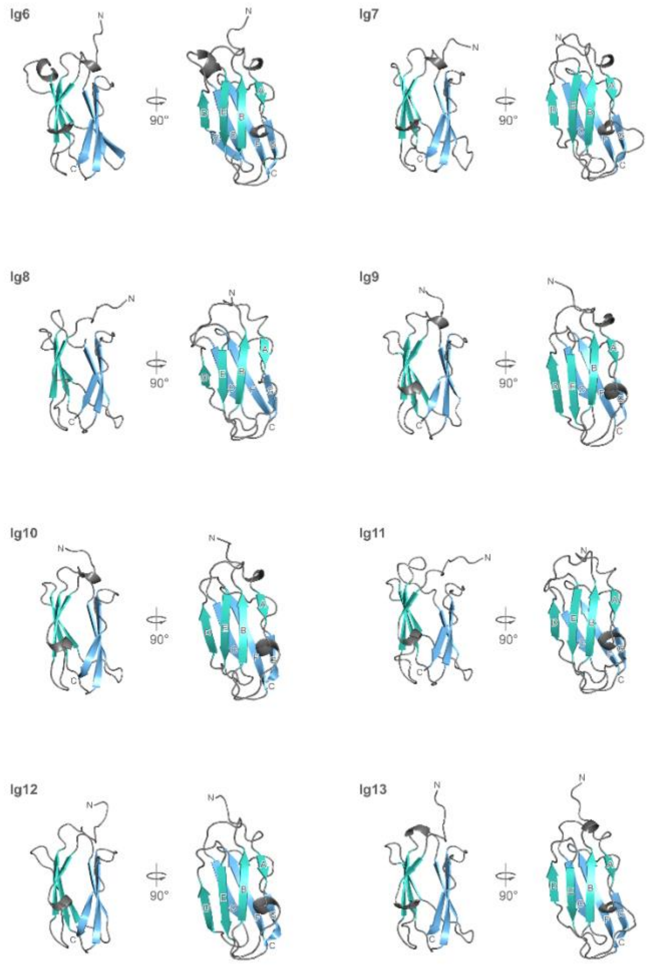

**Figure S1** (continue)

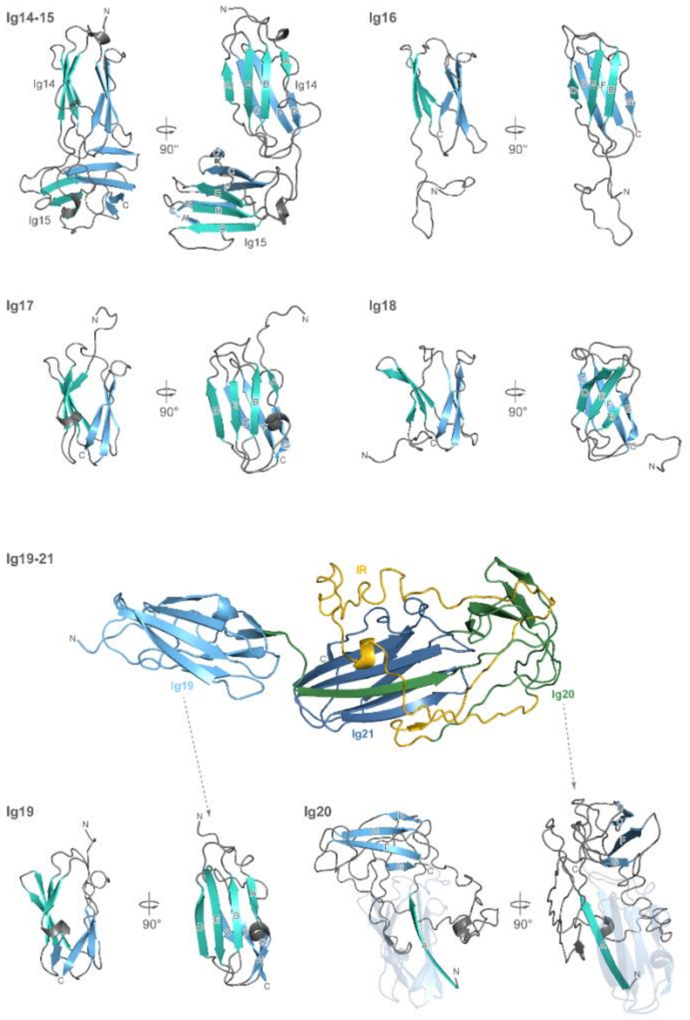

**Figure S1** (continue)

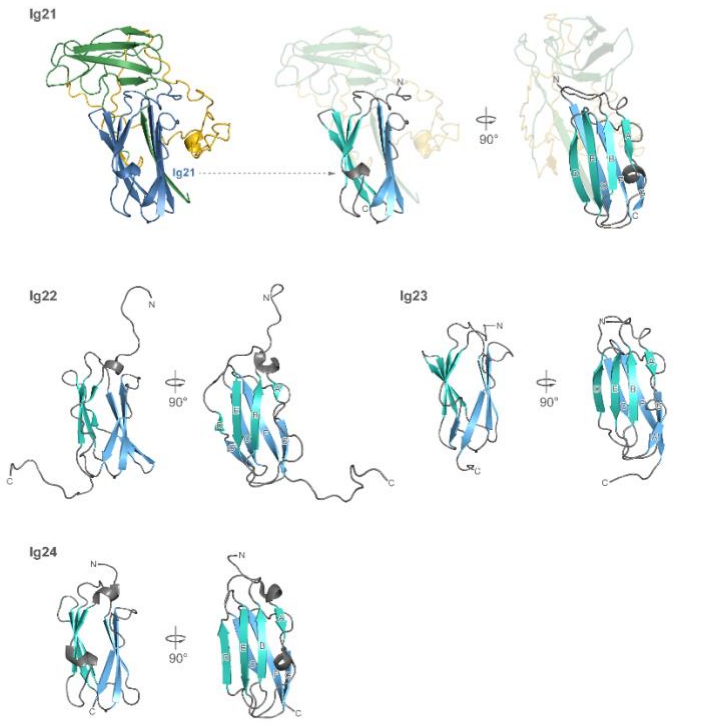

Figure S1

[Supplementary Figure S2](#)

Thermograms of individual Ig19, Ig21, Ig20-21 and Ig19-21 domains and constructs, respectively, showing relative order of unfolding.

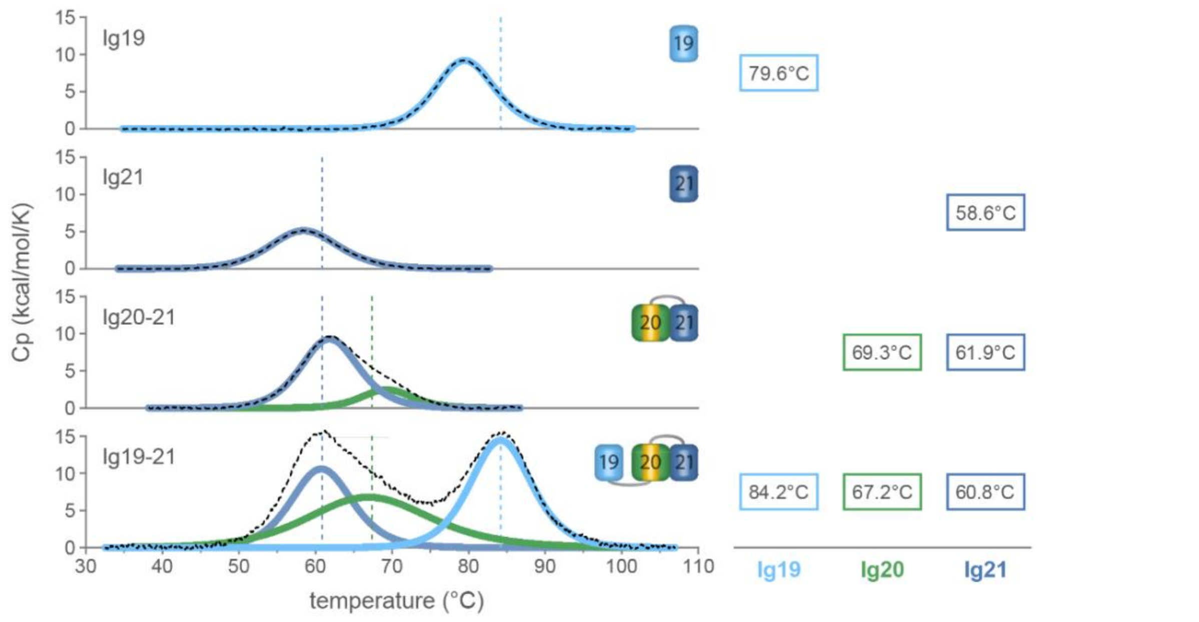

**Supplementary Figure S3**

Sequence alignment of human FlnC Ig domains showing higher conservation in deeper shades of red; the ß-strands names are as in topology notation (**Supplementary Figure S1 Structures + Topologies**), the length of the ß-strands is according to the consensus derived from superposition of experimentally determined structures (Ig4, Ig5, Ig14, Ig22, Ig23, Ig24). For clarity we omitted the insertion region (2162 - 2243) of Ig20, marked as *. PXSPF motif is boxed.

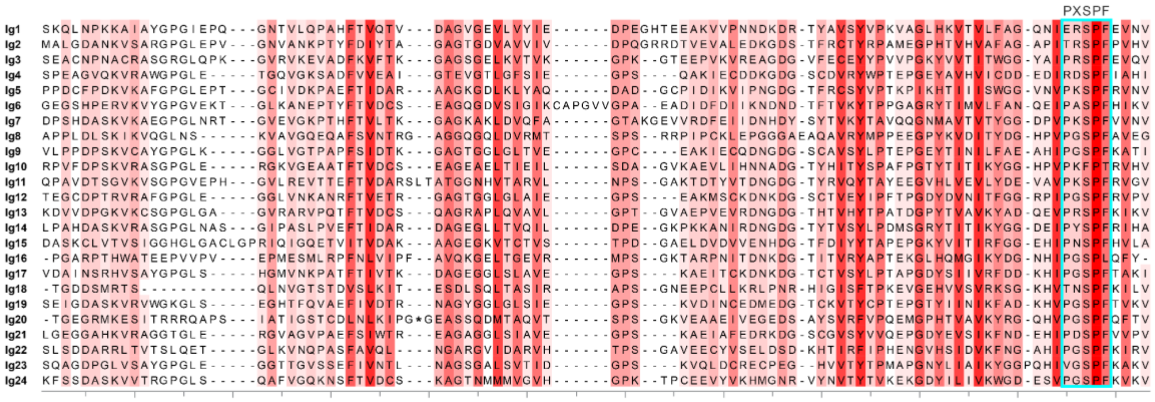

* Insertion region in FlnC: $^{2162}$GNWFQMVSAQERLTRTFTRSSHTYTRTERTEISKTRGGETKREVRVEESTQVGGDPFPAVFGDFLGRERLGSFGSITRQQEG$^{2243}$

Supplementary Figure S4

Ligplot representation of the interface between domains Ig14 and Ig15 observed in the crystal structures of the wild type construct and of the Ig14-15G1676R construct.

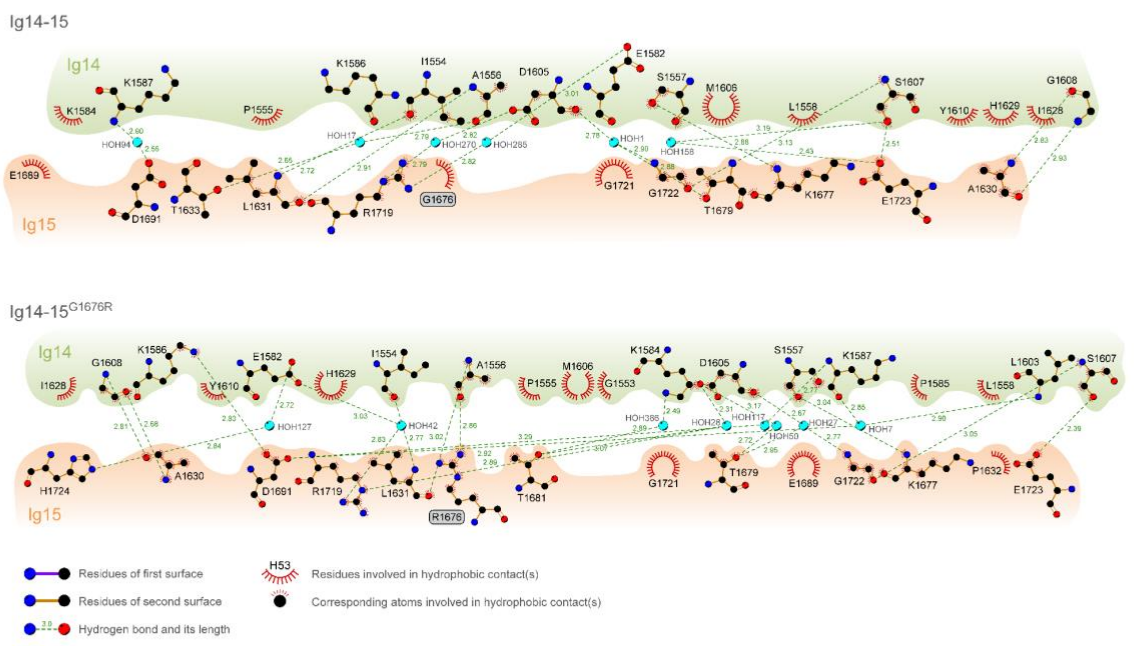

#### Supplementary Tables

**Table S1**.
Impact of selected variants on thermal stability analysed by DSC, and AMIVA-F prediction, and AMIVA-F prediction.

| **Domain** | **c-Notation** | **p-Notation** | **Phenotype** | $\Delta T_m$ (°C) | **Pathogenicity prediction (AMIVA-F)** | **Reference** |
|---|---|---|---|---|---|---|
| Ig14 | c.4871C>T | p.Ser1624Leu | **RCM** | Ig14 and Ig 15: -12.8 | **Yes** | This study and [6] |
| Ig15 | c.5026G>A | p.Gly1676Arg | **RCM** | Ig14: -6.5<br>Ig15: -23.9 [a] | **Yes** | This study |
| Ig19 | c.6173A>G | p.Gln2058Arg | *No* | Ig19: -2.8<br>Ig20: +0.3<br>Ig21: +0.4 | *No* | This study |
| Ig20 | c.6478A>T | p.Ile2160Phe | **RCM** | Ig19: -0.2<br>Ig20: +7.1<br>Ig21: -0.9 | **Yes** | [6] |
| Ig20 | c.6490T>C | p.Trp2164Cys | **HCM** | Ig19: -0.8<br>Ig20: +7.2<br>Ig21: +0.9 | **Yes** | H. Watkins, personal communication |
| Ig20 | c.6779A>G | p.Lys2260Arg | *No* | Ig19: -1.9<br>Ig20: +1.5<br>Ig21: +0.1 | *No* | This study |
| Ig20 | c.6892C>T | p.Pro2298Ser | **RCM** | Ig19: -3.6<br>Ig20: +6.9<br>Ig21: -0.7 | **Yes** | This study and [7] |

(a) Unfolding for Ig14-15 wildtype shows a single transition, this behaviour is lost upon introduction of p.Gly1676Arg.

**Table S2.**
Details about the crystal structure data.

| | Ig_d14-15 (7OUU) | Ig_14-15$^{S1624L}$ (7OUV) | Ig_14-15$^{G1676R}$ (7P0E) |
|---|---|---|---|
| **Data collection** | | | |
| Wavelength (Å) | 0,9686 | 0,9795 | 0,9655 |
| Resolution range (Å) | 19.83 - 1.47 (1.523 - 1.47)[a] | 37.97 - 1.8 (1.864 - 1.8)[a] | 38.26 - 1.6 (1.657 - 1.6)[a] |
| Space group | $P\ 2_1\ 2_1\ 2_1$ | $P\ 2_1\ 2_1\ 2_1$ | $P\ 2_1\ 2_1\ 2_1$ |
| Unit cell (Å) | 45.04 58.82 178.31 | 39.13 70.50 156.94 | 39.15 59.46 180.07 |
| Total reflections | 792556 (20899) | 520298 (54865) | 212835 (18674) |
| Unique reflections | 71856 (3680) | 41232 (4057) | 53624 (4752) |
| Multiplicity | 11.0 (5.7) | 12.6 (13.5) | 4.0 (3.9) |
| Completeness (%) | 88.0(45.8) | 99.6 (99.8) | 94.0 (79.6) |
| Mean *I*/sigma (*I*) | 11.1 (0.1) | 9.99 (0.8) | 9.97 (0.5) |
| Wilson B-factor (Å$^2$) | 24.1 | 33.7 | 29.8 |
| R-merge | 0.1136 | 0.1469 | 0.06401 |
| R-meas | 0.1187 | 0.1535 | 0.0733 |
| R-pim | 0.03397 | 0.04388 | 0.03494 |
| CC1/2 | 0.999 (0.019) | 0.998 (0.478) | 0.998 (0.134) |
| CC* | 1 (0.193) | 1 (0.804) | 0.999 (0.486) |
| **Refinement** | | | |
| Reflections used in refinement | 71848 (3680) | 41103 (4050) | 53270 (4458) |
| Reflections used for *R*-free | 1903 (97) | 1417 (140) | 1446 (121) |
| *R*-work | 0.1881 (0.3991) | 0.2134 (0.3352) | 0.1859 (0.3857) |
| *R*-free | 0.2277 (0.3750) | 0.2472 (0.3460) | 0.2211 (0.4270) |
| CC (work) | 0.955 (0.103) | 0.938 (0.722) | 0.964 (0.399) |
| CC (free) | 0.954 (0.096) | 0.940 (0.599) | 0.938 (0.285) |
| No. of non-hydrogen atoms | 3379 | 3376 | 3376 |
| No. of atoms macromolecules | 3015 | 3032 | 3028 |
| No. of atoms in ligands | 0 | 0 | 46 |
| No. of solvent atoms | 364 | 344 | 320 |
| Protein residues | 406 | 406 | 404 |
| RMS (bonds) (Å) | 0,002 | 0.004 | 0.015 |
| RMS (angles) (°) | 0.54 | 0.68 | 1.24 |
| Ramachandran favored (%) | 98.51 | 98.51 | 97.5 |
| Ramachandran allowed (%) | 1.49 | 1.24 | 2.5 |
| Ramachandran outliers (%) | 0 | 0.25 | 0 |
| Rotamer outliers (%) | 0 | 0.61 | 0 |
| Clashscore | 0.34 | 2.5 | 3.79 |
| Average B-factor (Å$^2$) | 35.0 | 42.4 | 38.9 |
| Macromolecules (Å$^2$) | 34.2 | 42.1 | 38.5 |
| Ligands (Å$^2$) | - | - | 50.7 |
| Solvent (Å$^2$) | 41.4 | 43.9 | 42.3 |
| Number of TLS groups | anisotropic B-factor model | 7 | 7 |

[a] Values in parentheses represent the highest resolution shell.

**Table S3.**
Table of human FlnC mutations reported in this work. Mutants experimentally characterized in this work are highlighted in bold.

| Domain | c-Notation | p-Notation | Phenotype | Reference |
|---|---|---|---|---|
| **ABD** | **c.245T>G** | **p.Met82Lys** | **DCM** | **This study** |
| ABD | c.318C>G | Phe106Leu | DCM | 8 |
| ABD | c.368T>C | Val123Ala | HCM | 7,9 |
| ABD | c.420C>A | His140Gln | HCM | 8 |
| ABD | c.433A>G | Met145Val | HCM | 10 |
| ABD | c.547C>T | Thr160Lys | ACM | 8 |
| ABD | c.577G>A | Arg183Cys | HCM | 8,10 |
| ABD | c.664A>G | Ala193Thr | Distal myopathy | 11,12 |
| ABD | c.664A>G | Met222Val | DM PM MFM | 13 |
| ABD | c.752T>C | Met251Thr | Distal myopathy | 11,12 |
| Ig1 | c.824C>T | Pro275Leu | HCM | 8 |
| Ig1 | c.870C>A | Asn290Lys | HCM | 14,15 |
| Ig1 | c.896C>T | Thr299Ile | HCM | 8 |
| Ig1 | c.925G>A | Glu309Lys | IBM, ALS, Dementia | 16 |
| Ig1 | c.987C>A | Asn329Lys | HCM | 17 |
| Ig1 | c.986A>G | Asn329Ser | HCM | 8,17 |
| Ig1 | c.1076T>C | Ile359Thr | HCM | 8 |
| Ig2 | c.1132G>T | Val378Leu | HCM | 8 |
| Ig2 | c.1322G>T | Arg441Ile | DCM | 18 |
| Ig4 | c.1882G>A | Val628Met | HCM | 10 |
| Ig4 | c.1942G>T | Asp648Tyr | MFM, DM | 19 |
| Ig5 | c.2050G>C | Val684Leu | HCM | 10 |
| Ig5 | c.2170G>A | Gly724Ser | HCM | 10 |
| Ig6 | c.2375G>T | Ser792Ile | HCM | 20 |
| Ig6 | c.2425G>A | Val809Met | DCM | 21 |
| Ig6 | c.2450T>C | Ile817Thr | HCM | 22 |
| Ig7 | c.2587C>T | Pro863Ser | HCM | 10 |
| Ig7 | c.2672C>T | Thr891Met | HCM | 8,10 |
| Ig7 | c.2737G>A | Glu913Lys | HCM | 8,10 |
| Ig7 | c.2839G>C | Gly947Arg | HCM | 8,10 |
| Ig8 | c.2888G>A | Pro963Arg | DCM | 18 |
| Ig8 | c.3092>X | Pro1031Leu | HCM | 8 |

| | | | | |
|---|---|---|---|---|
| Ig10 | c.3547_3548delinsCT | Ala1183Leu | RCM | [23] |
| Ig10 | c.3557C>T | Ala1186Val | PM, DM | [24] |
| Ig10 | c.3581C>T | Ser1194Leu | HCM | [25] |
| Ig10 | c.3592>X | Val1198Gly | DCM | [18] |
| Ig10 | c.3623C>T | Ala1208Val | HCM | [8,10] |
| Ig10 | c.3646T>A | Tyr1216Asn | HCM/MFM | [26] |
| Ig10 | c.3706C>T | Pro1236Ser | Muscular phenotype | [27] |
| Ig10 | c.3721C>T | Arg1241Cys | Inclusion body formation | [16] |
| Ig12 | c.4055G>A | Arg1352His | HCM | [8] |
| Ig12 | c.4222G>A | Glu1408Asp | HCM | [8] |
| Ig12 | c.4270G>A | Gly1424Val | HCM | [25] |
| Ig14 | c.4615G>A | Ala1539Thr | HCM | [14] |
| Ig14 | c.4636G>A | Gly1546Ser | RCM | [28] |
| Ig14 | c.4651G>A | Ala1551Thr | ALS, Dementia | [29] |
| Ig14 | c.4700G>A | Arg1567Gln | DCM | [30] |
| Ig14 | c.4795A>G | Thr1599Ala | HCM | [7,31] |
| **Ig14** | **c.4871C>T** | **Ser1624Leu** | **RCM** | **This study, [6]** |
| Ig14 | c.4916G>A | Cys1639Tyr | RCM | [18] |
| Ig15 | c.4997T>C | Ile1666Thr | NCCM | [25] |
| **Ig15** | **c.5026G>A** | **Gly1676Arg** | **RCM** | **This study** |
| Ig15 | c.5042C>T | Thr1681Met | HCM | [15] |
| Ig15 | c.5068C>T | Leu1690Phe | HCM | [15] |
| Ig15 | c.5071G>A | Asp1691Asn | MFM | [19] |
| Ig15 | c.5125C>T | Pro1709Ser | HCM | [10] |
| Ig15 | c.5132C>T | Pro1711Leu | HCM | [10] |
| Ig16 | c.5278G>A | Gly1760Ser | MFM | [27] |
| Ig16 | c.5524G>A | Asn1843Ser | HCM | [8] |
| Ig17 | c.5644A>G | Ile1882Val | ACM | [32] |
| Ig17 | c.5807A>C | His1936Pro | HCM | [8,10] |
| Ig18 | c.5888C>T | Thr1963Met | HCM | [10] |
| Ig18 | c.5954C>T | Ser1985Leu | HCM | [31] |
| Ig18 | c.5996G>A | Arg1999Gln | HCM | [20] |
| Ig18 | c.6032G>A | Gly2011Glu | HCM | [25] |
| Ig18 | c.6053G>A | Arg2018His | HCM | [33] |
| Ig19 | c.6115G>A | Gly2039Arg | HCM | [25] |

| | | | | |
|---|---|---|---|---|
| Ig19 | c.6134G>A | Arg2045Gln | HCM | [33] |
| Ig19 | c.6205G>A | Ala2069Thr | HCM | [31] |
| Ig19 | c.6208G>A | Gly2070Ser | RCM, ACM | [34] |
| Ig20 | c.6397C>T | Arg2133Cys | HCM | [10,35] |
| Ig20 | c.6398G>A | Arg2133His | HCM | [14,35] |
| Ig20 | c.6419G>A | Arg2140Gln | HCM | [15] |
| Ig20 | c.6451G>A | Gly2151Ser | HCM | [14] |
| **Ig20** | **c.6478A>T** | **Ile2160Phe** | **RCM** | **This study, [6]** |
| **IR** | c.6490T>C | **Trp2164Cys** | **HCM** | **This study** |
| IR | c.6538C>T | Arg2180Cys | ACM | [31] |
| IR | c.6559C>T | Arg2187Cys | ACM | [31] |
| IR | c.6589C>T | Arg2197Trp | HCM | [31] |
| Ig20 | c.6808G>A | Glu2270Lys | Muscular phenotype | [31] |
| Ig20 | c.6824G>T | Ser2275Ile | Muscular phenotype | [31] |
| Ig20 | c.6889G>A | Val2297Met | RCM | [36] |
| **Ig20** | **c.6892C>T** | **Pro2298Ser** | **RCM** | **This study, [37]** |
| **Ig20** | **c.6892C>T** | **Pro2298Leu** | **HCM** | **This study** |
| Ig20 | c.6895G>A | Gly2299Ser | HCM | [25] |
| Ig20 | c.6902C>T | Pro2301Leu | HCM | [38] |
| Ig20 | c.6901C>G | Pro2301Ala | HCM | [15] |
| Ig20 | c.6943C>A | His2315Asn | HCM | [14] |
| Ig21 | c.6952C>T | Arg2318Trp | HCM | [15] |
| Ig21 | c.7030G>A | Ala2344Thr | HCM | [8,10] |
| Ig21 | c.7034G>A | Gly2345Glu | Congenital heart disease | [39] |
| Ig21 | c.7076T>C | Ile2359Thr | HCM | [25] |
| Ig21 | c.7123G>T | Val2375Phe | HCM | [15] |
| Ig21 | c.7123G>C | Val2375Leu | HCM | [25] |
| Ig21 | c.7123G>A | Val2375Ile | MCM | [8] |
| Ig21 | c.7177C>T | Pro2393Ser | Noncompaction cardiomyopathy | [31,35] |
| Ig21 | c.7186C>A | Pro2396Thr | HCM | [10] |
| Ig22 | c.7228C>T | Arg2410Cys | HCM | [25] |
| Ig22 | c.7250A>C | Gln2417Pro | HCM | [25] |
| Ig22 | c.7256C>T | Thr2419Met | MFM | [40] |
| Ig22 | c.7289C>T | Ala2430Val | HCM | [14,41] |
| Ig22 | c.7484G>A | Arg2495His | HCM | [42] |

| | | | | |
|---|---|---|---|---|
| Ig23 | c.7514C>T | Pro2505Leu | HCM | [10] |
| Ig23 | c.7688A>G | Tyr2563Cys | RCM | [37] |
| Ig23 | c.7724T>A | Ile2575Asn | PM, CON | [31] |
| Ig23 | c.7781G>C | Gly2594Ala | HCM | [33] |
| Ig24 | c.7942G>T | Ala2648Ser | Arrhythmia | [18] |

| Table S4. Mutations selected to represent the non-pathogenic dataset in AMIVA-F. All mutations were selected from GnomAD database and fulfil the criteria outlined in **Selection of training sets**. | | | | | | | |
|---|---|---|---|---|---|---|---|
| **Domain** | **Chromosome** | **c-Notation** | **p-Notation** | **Allele Count** | **Allele Number** | **Allele Frequency** | **Homozygote Count** |
| ABD | 7 | c.22T>C | p.Ser8Pro | 31 | 275298 | 0.00011261 | 0 |
| Ig1 | 7 | c.1081C>T | p.Arg361Cys | 49 | 280936 | 0.00017442 | 0 |
| Ig2 | 7 | c.1108A>G | p.Met370Val | 31 | 280940 | 0.00011034 | 0 |
| | 7 | c.1142G>A | p.Arg381His | 31 | 280900 | 0.00011036 | 0 |
| | 7 | c.1147C>G | p.Pro383Ala | 31 | 249528 | 0.00012424 | 0 |
| | 7 | c.1166G>A | p.Gly389Asp | 35 | 280866 | 0.00012462 | 0 |
| | 7 | c.1354G>A | p.Val452Met | 38 | 279848 | 0.00013579 | 0 |
| Ig3 | 7 | c.1474A>G | p.Lys492Glu | 73 | 280852 | 0.00025992 | 0 |
| | 7 | c.1519G>A | p.Gly507Arg | 221 | 280698 | 0.00078732 | 0 |
| | 7 | c.1576C>T | p.Arg526Trp | 46 | 280892 | 0.00016376 | **1** |
| | 7 | c.1577G>A | p.Arg526Gln | 614 | 280918 | 0.00218569 | **4** |
| | 7 | c.1600G>A | p.Glu534Lys | 235 | 280962 | 0.00083641 | 0 |
| | 7 | c.1657G>A | p.Gly553Ser | 76 | 280938 | 0.00027052 | 0 |
| Ig4 | 7 | c.1976T>G | p.Leu659Arg | 102 | 280798 | 0.00036325 | **1** |
| Ig5 | 7 | c.2068T>C | p.Phe690Leu | 48 | 280970 | 0.00017084 | 0 |
| | 7 | c.2078A>C | p.Asp693Ala | 967 | 280970 | 0.00344165 | **2** |
| | 7 | c.2125G>A | p.Ala709Thr | 295 | 279720 | 0.00105463 | **1** |
| | 7 | c.2128G>A | p.Asp710Asn | 86 | 279874 | 0.00030728 | 0 |
| | 7 | c.2180G>A | p.Arg727His | 108 | 280538 | 0.00038498 | 0 |
| Ig6 | 7 | c.2296C>T | p.Arg766Trp | 21 | 183654 | 0.00011435 | 0 |
| | 7 | c.2392G>A | p.Asp798Asn | 63 | 279436 | 0.00022545 | **1** |
| | 7 | c.2501C>T | p.Thr834Met | 2083 | 280520 | 0.0074255 | **62** |
| | 7 | c.2507C>A | p.Pro836Gln | 193 | 280496 | 0.00068807 | **0** |
| Ig7 | 7 | c.2635C>T | p.Arg879Cys | 39 | 277130 | 0.00014073 | 0 |

| | | | | | | | |
|---|---|---|---|---|---|---|---|
| | 7 | c.2686G>A | p.Gly896Arg | 110 | 280648 | 0.00039195 | 0 |
| | 7 | c.2839G>C | p.Gly947Arg | 45 | 280820 | 0.00016025 | 0 |
| Ig8 | 7 | c.3145G>T | p.Gly1049Cys | 43 | 278222 | 0.00015455 | 0 |
| | 7 | c.3146G>T | p.Gly1049Val | 43 | 278172 | 0.00015458 | 0 |
| Ig9 | 7 | c.3242C>T | p.Ala1081Val | 36 | 280412 | 0.00012838 | 0 |
| | 7 | c.3304C>T | p.Pro1102Ser | 30 | 280776 | 0.00010685 | 0 |
| Ig10 | 7 | c.3506A>G | p.Lys1169Arg | 283 | 280580 | 0.00100863 | **2** |
| | 7 | c.3721C>T | p.Arg1241Cys | 1729 | 279050 | 0.00619602 | **17** |
| Ig11 | 7 | c.3757G>A | p.Val1253Ile | 365 | 278306 | 0.00131151 | **5** |
| | 7 | c.3812C>G | p.Thr1271Ser | 46 | 248868 | 0.00018484 | 0 |
| | 7 | c.3847A>G | p.Thr1283Ala | 32 | 280250 | 0.00011418 | 0 |
| | 7 | c.3938G>A | p.Arg1313Gln | 57 | 280412 | 0.00020327 | 0 |
| | 7 | c.4022G>A | p.Arg1341Gln | 264 | 280120 | 0.00094245 | **1** |
| Ig12 | 7 | c.4097A>G | p.Asn1366Ser | 44 | 273358 | 0.00016096 | 0 |
| Ig13 | 7 | c.4459G>A | p.Val1487Met | 60 | 278014 | 0.00021582 | **1** |
| | 7 | c.4553A>G | p.Lys1518Arg | 52 | 264562 | 0.00019655 | 0 |
| Ig14 | 7 | c.4651G>A | p.Ala1551Thr | 33 | 280162 | 0.00011779 | 0 |
| | 7 | c.4700G>A | p.Arg1567Gln | 20774 | 279986 | 0.07419657 | **872** |
| | 7 | c.4763C>G | p.Ala1588Gly | 45 | 280652 | 0.00016034 | 0 |
| Ig15 | 7 | c.4970G>A | p.Arg1657Gln | 29 | 280592 | 0.00010335 | 0 |
| | 7 | c.5020G>A | p.Gly1674Ser | 38 | 280830 | 0.00013531 | 0 |
| | 7 | c.5042C>G | p.Thr1681Arg | 199 | 280908 | 0.00070842 | 0 |
| | 7 | c.5143G>A | p.Val1715Ile | 62 | 280826 | 0.00022078 | 0 |
| Ig16 | 7 | c.5278G>A | p.Gly1760Ser | 29 | 176390 | 0.00016441 | 0 |
| | 7 | c.5284C>T | p.Arg1762Cys | 44 | 175150 | 0.00025121 | 0 |
| | 7 | c.5311C>G | p.Pro1771Ala | 71 | 280832 | 0.00025282 | 0 |

| | | | | | | | |
|---|---|---|---|---|---|---|---|
| | 7 | c.5374G>A | p.Ala1792Thr | 68 | 280634 | 0.00024231 | 0 |
| | 7 | c.5375C>T | p.Ala1792Val | 27 | 249226 | 0.00010834 | 0 |
| Ig17 | 7 | c.5578C>T | p.Arg1860Cys | 1463 | 280876 | 0.0052087 | **9** |
| | 7 | c.5644A>G | p.Ile1882Val | 312 | 280886 | 0.00111077 | **1** |
| | 7 | c.5764G>A | p.Ala1922Thr | 415 | 282370 | 0.0014697 | **4** |
| Ig18 | 7 | c.5944C>T | p.Arg1982Cys | 45 | 280056 | 0.00016068 | 0 |
| | 7 | c.5954C>T | p.Ser1985Leu | 56 | 280118 | 0.00019992 | 0 |
| Ig19 | 7 | c.6175G>A | p.Val2059Met | 238 | 279776 | 0.00085068 | 0 |
| Ig20 | 7 | c.6808G>A | p.Glu2270Lys | 193 | 260954 | 0.00073959 | 0 |
| | 7 | c.6865G>A | p.Ala2289Thr | 35 | 265342 | 0.00013191 | 0 |
| Ig21 | 7 | c.6988G>A | p.Gly2330Ser | 166 | 277392 | 0.00059843 | **2** |
| | 7 | c.6991G>A | p.Val2331Met | 129 | 277812 | 0.00046434 | 0 |
| | 7 | c.7091G>A | p.Arg2364His | 484 | 280776 | 0.00172379 | **1** |
| Ig22 | 7 | c.7256C>T | p.Thr2419Met | 42 | 245700 | 0.00017094 | 0 |
| | 7 | c.7289C>T | p.Ala2430Val | 28 | 278082 | 0.00010069 | 0 |
| | 7 | c.7291G>A | p.Val2431Met | 47 | 246748 | 0.00019048 | 0 |
| Ig23 | 7 | c.7614G>T | p.Leu2538Phe | 435 | 279574 | 0.00155594 | 3 |
| Ig24 | 7 | c.8003T>C | p.Met2668Thr | 219 | 262788 | 0.00083337 | 0 |

**Table S5.**
Experimental structures of FlnC domains, determined by X-ray crystallography or by NMR, and experimental structures of FlnA and FlnB domains used as templates to generate homology models, with Modeller, of FlnC domains. The resolution of the X-ray crystal structures and the percentage of sequence identity between template and target are provided.

| Domain | Position | Template (pdb id/protein name) | Resolution (Å) | Seq. identity (%) |
|---|---|---|---|---|
| ABD | 36-262 | 3hop/FlnA | 2.3 | 85.8 |
| Ig1 | 270-368 | 4b7l/FlnB | 2.1 | 67.7 |
| Ig2 | 370-468 | 4b7l/FlnB | 2.1 | 37.4 |
| Ig3 | 469-565 | 6ew1/FlnA | 2.3 | 77.3 |
| Ig4-5 | 568-760 | 3v8o/FlnC | 2.8 | na[b] |
| Ig6 | 759-861 | 4m9p/FlnA | 1.7 | 38.8 |
| Ig7 | 862-960 | 4m9p/FlnA | 1.7 | 40.4 |
| Ig8 | 961-1056 | 2dic/FlnB | na[a] | 41.7 |
| Ig9 | 1057-1149 | 2di9/FlnB | na[a] | 82.8 |
| Ig10 | 1150-1244 | 3rgh/FlnA | 2.4 | 61.1 |
| Ig11 | 1245-1344 | 2dib/FlnA | na[a] | 60 |
| Ig12 | 1345-1437 | 2dic/FlnB | na[a] | 69.9 |
| Ig13 | 1438-1533 | 2dj4/FlnB | na[a] | 71.9 |
| Ig14-15 | 1535-1758 | Crystal structure reported here | 1.8 | na[b] |
| Ig16 | 1759-1853 | 2d7n/FlnC | na[a] | na[b] |
| Ig17 | 1854-1946 | 2d7o/FlnC | na[a] | na[b] |
| Ig18 | 1947-2033 | 2k7q/FlnA | na[a] | na[b] |
| Ig19-21* | 2036-2401 | 2j3s/FlnA | 2.5 | 81.4* |
| Ig22 | 2398-2509 | 2D7P/FlnC | na[a] | na[b] |
| Ig23 | 2502-2598 | 2NQC/FlnC | 2.1 | na[b] |
| Ig24 | 2633-2725 | 1V05/FlnC | 1.4 | na[b] |

[a] If no resolution is given, the structure was determined by NMR.

[b] The percentage of sequence identity between target and template is not provided for experimental structures determined either with X-ray crystallography or NMR.

* The model was generated *via* i-Tasser [43].

**Table S6.**
Pathogenicity prediction of 16 mutations absent from the learning sets used for neural network training, which are rarely observed in the gnomeAD database, and for which the pathogenicity is unknown. Pathogenic predictions are indicated in **bold**, neutral in *italics*.

| **Domain** | **c-Notation** | **p-Notation** | **Pathogenicity prediction (AMIVA-F)** |
|---|---|---|---|
| ABD | c.431C>T | **Ser144Phe** | **Yes** |
| Ig1 | c.1019A>G | **Tyr340Cys** | **Yes** |
| Ig1 | c.1027A>G | *Lys343Glu* | *No* |
| Ig3 | c.1649C>A/G/T | *Thr550Met* | *No* |
| Ig5 | c.2272G>A | **Val758Met** | **Yes** |
| Ig8 | 2911G>A/T | **Val971Ile** | **Yes** |
| Ig11 | c.3742G>A/T | *Val1248Phe* | *No* |
| Ig11 | c.3905C>T | *Thr1302Ile* | *No* |
| Ig14 | c.5036C>A | **Thr1679Lys** | **Yes** |
| Ig15 | c.5205T>G | **Thr1735Trp** | **Yes** |
| Ig16 | c.5333G>A | **Arg1811Gln** | **Yes** |
| Ig17 | c.5764G>A/T | *Pro1920Ser* | *No* |
| Ig19 | c.6133C>A/T | **Arg2045Trp** | **Yes** |
| Ig20 | c.6808G>A/C | *Glu2270Gln* | *No* |
| Ig20 | c.6809A>C/T | *Glu2270Ala* | *No* |
| Ig22 | c.7302C>G/T | **Asn2434Lys** | **Yes** |

**Table S7.**
Comparison of predictors using 173 variants used in the learning set

| | AMIVA-F | Polyphen2 | SIFT | Provean |
|---|---|---|---|---|
| Correct Predictions (TP + TN) | 136 | 127 | 111 | 114 |
| False Predictions (FP + FN) | 37 | 46 | 62 | 59 |
| Accuracy | 78.6% | 73% | 64.2% | 65.9% |

[a] TP = true positives: number of pathogenic mutations that are predicted to be pathogenic.
[b] FN = false negatives: number of pathogenic mutations that are predicted to be non-pathogenic.
[d] FP = false positives: number of non-pathogenic mutations that are predicted to be pathogenic.
[c] TN = true negatives: number of non-pathogenic mutations that are predicted to be non-pathogenic.

**Table S8**.
Technical details on the algorithm used to predict human FlnC variants pathogenicity.

| | |
|---|---|
| Algorithm | Multilayer perceptron |
| Batch size | 100 |
| Hidden layers | a,2,3 |
| Learning rate | 0.78 |
| Momentum | 0.2 |
| Epochs | 517 |
| Threshold function | sigmoid |
| Validation threshold | 20 |
| Seed | 0 |
| Filters | Nominal to binary, normalized attributes, normalized numeric classes |

**Table S9**.
Pathogenicity propensities upon mutation of the 20 amino acids

| Amino acid type | Propensity (equation 1, Supplementary Information) |
|---|---|
| R | 3.666 |
| M | 3.058 |
| P | 1.584 |
| W | 1.402 |
| I | 1.318 |
| A | 1.294 |
| N | 1.216 |
| H | 1.183 |
| G | 1.086 |
| T | 1.062 |
| V | 0.924 |
| S | 0.751 |
| Q | 0.655 |
| E | 0.573 |
| C | 0.561 |
| D | 0.321 |
| Y | 0.315 |
| F | 0.274 |
| L | 0.180 |
| K | 0.000 |

#### References

1. Iwaoka, M., Katsuda, T., Komatsu, H., and Tomoda, S. (2005). Experimental and theoretical studies on the nature of weak nonbonded interactions between divalent selenium and halogen atoms. J Org Chem *70*, 321-327. 10.1021/jo048436a.
2. Wu, Q., Schapira, M., Arrowsmith, C.H., and Barsyte-Lovejoy, D. (2021). Protein arginine methylation: from enigmatic functions to therapeutic targeting. Nat Rev Drug Discov *20*, 509-530. 10.1038/s41573-021-00159-8.
3. Kozlowski, L.P., and Bujnicki, J.M. (2012). MetaDisorder: a meta-server for the prediction of intrinsic disorder in proteins. BMC Bioinformatics *13*, 111. 10.1186/1471-2105-13-111.
4. Eudes, R., Le Tuan, K., Delettre, J., Mornon, J.P., and Callebaut, I. (2007). A generalized analysis of hydrophobic and loop clusters within globular protein sequences. BMC Struct Biol *7*, 2. 10.1186/1472-6807-7-2.
5. Banani, S.F., Lee, H.O., Hyman, A.A., and Rosen, M.K. (2017). Biomolecular condensates: organizers of cellular biochemistry. Nature Reviews Molecular Cell Biology *18*, 285-298. 10.1038/nrm.2017.7.
6. Brodehl, A., Ferrier, R.A., Hamilton, S.J., Greenway, S.C., Brundler, M.-A., Yu, W., Gibson, W.T., McKinnon, M.L., McGillivray, B., Alvarez, N., et al. (2016). Mutations in FLNC are Associated with Familial Restrictive Cardiomyopathy. Human Mutation *37*, 269-279. https://doi.org/10.1002/humu.22942.
7. Gómez, J., Lorca, R., Reguero, J.R., Morís, C., Martín, M., Tranche, S., Alonso, B., Iglesias, S., Alvarez, V., Díaz-Molina, B., Avanzas, P., and Coto, E. (2017). Screening of the Filamin C Gene in a Large Cohort of Hypertrophic Cardiomyopathy Patients. Circulation: Cardiovascular Genetics *10*, e001584. 10.1161/circgenetics.116.001584.
8. Eden, M., and Frey, N. (2021). Cardiac Filaminopathies: Illuminating the Divergent Role of Filamin C Mutations in Human Cardiomyopathy. Journal of Clinical Medicine *10*, 577. 10.3390/jcm10040577.
9. Brodehl, A., Ferrier, R.A., Hamilton, S.J., Greenway, S.C., Brundler, M.-A., Yu, W., Gibson, W.T., McKinnon, M.L., McGillivray, B., Alvarez, N., et al. (2016). Mutations in<i>FLNC</i>are Associated with Familial Restrictive Cardiomyopathy. Human Mutation *37*, 269-279. 10.1002/humu.22942.
10. Cui, H., Wang, J., Zhang, C., Wu, G., Zhu, C., Tang, B., Zou, Y., Huang, X., Hui, R., Song, L., and Wang, S. (2018). Mutation profile of FLNC gene and its prognostic relevance in patients with hypertrophic cardiomyopathy. Molecular Genetics & Genomic Medicine *6*, 1104-1113. https://doi.org/10.1002/mgg3.488.
11. Duff, R.M., Tay, V., Hackman, P., Ravenscroft, G., McLean, C., Kennedy, P., Steinbach, A., Schöffler, W., van der Ven, P.F.M., Fürst, D.O., et al. (2011). Mutations in the N-terminal actin-binding domain of filamin C cause a distal myopathy. American journal of human genetics *88*, 729-740. 10.1016/j.ajhg.2011.04.021.
12. Wadmore, K., Azad, A.J., and Gehmlich, K. (2021). The Role of Z-disc Proteins in Myopathy and Cardiomyopathy. International Journal of Molecular Sciences *22*, 3058. 10.3390/ijms22063058.
13. Gemelli, C., Prada, V., Fiorillo, C., Fabbri, S., Maggi, L., Geroldi, A., Gibertini, S., Mandich, P., Trevisan, L., Fossa, P., et al. (2019). A novel mutation in the N-terminal acting-binding domain of Filamin C protein causing a distal myofibrillar myopathy. Journal of the Neurological Sciences *398*, 75-78. https://doi.org/10.1016/j.jns.2019.01.019.
14. Valdes-Mas, R., Gutierrez-Fernandez, A., Gomez, J., Coto, E., Astudillo, A., Puente, D.A., Reguero, J.R., Alvarez, V., Moris, C., Leon, D., et al. (2014). Mutations in filamin C cause a new form of familial hypertrophic cardiomyopathy. Nat Commun *5*, 5326. 10.1038/ncomms6326.
15. Gómez, J., Lorca, R., Reguero Julian, R., Morís, C., Martín, M., Tranche, S., Alonso, B., Iglesias, S., Alvarez, V., Díaz-Molina, B., Avanzas, P., and Coto, E. (2017). Screening of the Filamin C Gene in a Large Cohort of Hypertrophic Cardiomyopathy Patients. Circulation: Cardiovascular Genetics *10*, e001584. 10.1161/circgenetics.116.001584.

16. Weihl, C.C., Baloh, R.H., Lee, Y., Chou, T.-F., Pittman, S.K., Lopate, G., Allred, P., Jockel-Balsarotti, J., Pestronk, A., and Harms, M.B. (2015). Targeted sequencing and identification of genetic variants in sporadic inclusion body myositis. Neuromuscular Disorders *25*, 289-296. https://doi.org/10.1016/j.nmd.2014.12.009.
17. Alejandra Restrepo-Cordoba, M., Campuzano, O., Ripoll-Vera, T., Cobo-Marcos, M., Mademont-Soler, I., Gámez, J.M., Dominguez, F., Gonzalez-Lopez, E., Padron-Barthe, L., Lara-Pezzi, E., et al. (2017). Usefulness of Genetic Testing in Hypertrophic Cardiomyopathy: an Analysis Using Real-World Data. Journal of Cardiovascular Translational Research *10*, 35-46. 10.1007/s12265-017-9730-8.
18. Xiao, F., Wei, Q., Wu, B., Liu, X., Mading, A., Yang, L., Li, Y., Liu, F., Pan, X., and Wang, H. (2020). Clinical exome sequencing revealed that FLNC variants contribute to the early diagnosis of cardiomyopathies in infant patients. Transl Pediatr *9*, 21-33. 10.21037/tp.2019.12.02.
19. Zhang, Y.-T., Pu, C.-Q., Ban, R., Liu, H.-X., Shi, Q., and Lu, X.-H. (2018). Clinical, Pathological, and Genetic Features of Two Chinese Cases with Filamin C Myopathy. Chinese Medical Journal *131*.
20. Jaafar, N., Gómez, J., Kammoun, I., Zairi, I., Amara, W.B., Kachboura, S., Kraiem, S., Hammami, M., Iglesias, S., Alonso, B., and Coto, E. (2016). Spectrum of Mutations in Hypertrophic Cardiomyopathy Genes Among Tunisian Patients. Genetic Testing and Molecular Biomarkers *20*, 674-679. 10.1089/gtmb.2016.0187.
21. Janin, A., N'Guyen, K., Habib, G., Dauphin, C., Chanavat, V., Bouvagnet, P., Eschalier, R., Streichenberger, N., Chevalier, P., and Millat, G. (2017). Truncating mutations on myofibrillar myopathies causing genes as prevalent molecular explanations on patients with dilated cardiomyopathy. Clinical Genetics *92*, 616-623. 10.1111/cge.13043.
22. Cirino Allison, L., Lakdawala Neal, K., McDonough, B., Conner, L., Adler, D., Weinfeld, M., O'Gara, P., Rehm Heidi, L., Machini, K., Lebo, M., et al. (2017). A Comparison of Whole Genome Sequencing to Multigene Panel Testing in Hypertrophic Cardiomyopathy Patients. Circulation: Cardiovascular Genetics *10*, e001768. 10.1161/circgenetics.117.001768.
23. Kiselev, A., Vaz, R., Knyazeva, A., Khudiakov, A., Tarnovskaya, S., Liu, J., Sergushichev, A., Kazakov, S., Frishman, D., Smolina, N., et al. (2018). De novo mutations in FLNC leading to early-onset restrictive cardiomyopathy and congenital myopathy. Human Mutation *39*, 1161-1172. https://doi.org/10.1002/humu.23559.
24. Ghaoui, R., Cooper, S.T., Lek, M., Jones, K., Corbett, A., Reddel, S.W., Needham, M., Liang, C., Waddell, L.B., Nicholson, G., et al. (2015). Use of Whole-Exome Sequencing for Diagnosis of Limb-Girdle Muscular Dystrophy: Outcomes and Lessons Learned. JAMA Neurology *72*, 1424-1432. 10.1001/jamaneurol.2015.2274.
25. Ader, F., De Groote, P., Réant, P., Rooryck-Thambo, C., Dupin-Deguine, D., Rambaud, C., Khraiche, D., Perret, C., Pruny, J.F., Mathieu-Dramard, M., et al. (2019). FLNC pathogenic variants in patients with cardiomyopathies: Prevalence and genotype-phenotype correlations. Clinical Genetics *96*, 317-329. https://doi.org/10.1111/cge.13594.
26. Avila-Smirnow, D., Béhin, A., Gueneau, L., Claeys, K., Beuvin, M., Goudeau, B., Richard, P., Yaou, R.B., Romero, N.B., Mathis, S., et al. (2010). P2.18 A novel missense FLNC mutation causes arrhythmia and late onset myofibrillar myopathy with particular histopathology features. Neuromuscular Disorders *20*, 623-624. https://doi.org/10.1016/j.nmd.2010.07.090.
27. Yu, M., Zheng, Y., Jin, S., Gang, Q., Wang, Q., Yu, P., Lv, H., Zhang, W., Yuan, Y., and Wang, Z. (2017). Mutational spectrum of Chinese LGMD patients by targeted next-generation sequencing. PLOS ONE *12*, e0175343. 10.1371/journal.pone.0175343.
28. Sanoja, A.J., Li, H., Fricker, F.J., Kingsmore, S.F., and Wallace, M.R. (2018). Exome sequencing identifies <i>FLNC</i> and <i>ADD3</i> variants in a family with cardiomyopathy. Journal of Translational Genetics and Genomics *2*, 6. 10.20517/jtgg.2017.13.
29. Janssens, J., Philtjens, S., Kleinberger, G., Van Mossevelde, S., van der Zee, J., Cacace, R., Engelborghs, S., Sieben, A., Banzhaf-Strathmann, J., Dillen, L., et al. (2015). Investigating the

role of filamin C in Belgian patients with frontotemporal dementia linked to GRN deficiency in FTLD-TDP brains. Acta Neuropathol Commun *3*, 68-68. 10.1186/s40478-015-0246-7.

30. Esslinger, U., Garnier, S., Korniat, A., Proust, C., Kararigas, G., Müller-Nurasyid, M., Empana, J.-P., Morley, M.P., Perret, C., Stark, K., et al. (2017). Exome-wide association study reveals novel susceptibility genes to sporadic dilated cardiomyopathy. PLOS ONE *12*, e0172995. 10.1371/journal.pone.0172995.
31. Verdonschot, J.A.J., Vanhoutte, E.K., Claes, G.R.F., Helderman-van den Enden, A., Hoeijmakers, J.G.J., Hellebrekers, D., de Haan, A., Christiaans, I., Lekanne Deprez, R.H., Boen, H.M., et al. (2020). A mutation update for the FLNC gene in myopathies and cardiomyopathies. Hum Mutat *41*, 1091-1111. 10.1002/humu.24004.
32. Hall, C.L., Akhtar, M.M., Sabater-Molina, M., Futema, M., Asimaki, A., Protonotarios, A., Dalageorgou, C., Pittman, A.M., Suarez, M.P., Aguilera, B., et al. (2020). Filamin C variants are associated with a distinctive clinical and immunohistochemical arrhythmogenic cardiomyopathy phenotype. International Journal of Cardiology *307*, 101-108. 10.1016/j.ijcard.2019.09.048.
33. Chanavat, V., Janin, A., and Millat, G. (2016). A fast and cost-effective molecular diagnostic tool for genetic diseases involved in sudden cardiac death. Clinica Chimica Acta *453*, 80-85. 10.1016/j.cca.2015.12.011.
34. Ortiz-Genga, M.F., Cuenca, S., Dal Ferro, M., Zorio, E., Salgado-Aranda, R., Climent, V., Padrón-Barthe, L., Duro-Aguado, I., Jiménez-Jáimez, J., Hidalgo-Olivares, V.M., et al. (2016). Truncating FLNC Mutations Are Associated With High-Risk Dilated and Arrhythmogenic Cardiomyopathies. Journal of the American College of Cardiology *68*, 2440-2451. 10.1016/j.jacc.2016.09.927.
35. van Waning, J.I., Hoedemaekers, Y.M., te Rijdt, W.P., Jpma, A.I., Heijsman, D., Caliskan, K., Hoendermis, E.S., Willems, T.P., Wijngaard, A.v.d., Suurmeijer, A., et al. (2019). FLNC missense variants in familial noncompaction cardiomyopathy. Cardiogenetics *9*. 10.4081/cardiogenetics.2019.8181.
36. Tucker, N.R., McLellan, M.A., Hu, D., Ye, J., Parsons, V.A., Mills, R.W., Clauss, S., Dolmatova, E., Shea, M.A., Milan, D.J., et al. (2017). Novel Mutation in FLNC (Filamin C) Causes Familial Restrictive Cardiomyopathy. Circ Cardiovasc Genet *10*. 10.1161/CIRCGENETICS.117.001780.
37. Schubert, J., Tariq, M., Geddes, G., Kindel, S., Miller, E.M., and Ware, S.M. (2018). Novel pathogenic variants in filamin C identified in pediatric restrictive cardiomyopathy. Hum Mutat *39*, 2083-2096. 10.1002/humu.23661.
38. Roldán-Sevilla, A., Palomino-Doza, J., de Juan, J., Sánchez, V., Domínguez-González, C., Salguero-Bodes, R., and Arribas-Ynsaurriaga, F. (2019). Missense Mutations in the FLNC Gene Causing Familial Restrictive Cardiomyopathy. Circulation: Genomic and Precision Medicine *12*, e002388. 10.1161/circgen.118.002388.
39. Kosmicki, J.A., Samocha, K.E., Howrigan, D.P., Sanders, S.J., Slowikowski, K., Lek, M., Karczewski, K.J., Cutler, D.J., Devlin, B., Roeder, K., et al. (2017). Refining the role of de novo protein-truncating variants in neurodevelopmental disorders by using population reference samples. Nature Genetics *49*, 504-510. 10.1038/ng.3789.
40. Tasca, G., Odgerel, Z., Monforte, M., Aurino, S., Clarke, N.F., Waddell, L.B., Udd, B., Ricci, E., and Goldfarb, L.G. (2012). Novel FLNC mutation in a patient with myofibrillar myopathy in combination with late-onset cerebellar ataxia. Muscle & Nerve *46*, 275-282. https://doi.org/10.1002/mus.23349.
41. Schänzer, A., Schumann, E., Zengeler, D., Gulatz, L., Maroli, G., Ahting, U., Sprengel, A., Gräf, S., Hahn, A., Jux, C., et al. (2021). The p.Ala2430Val mutation in filamin C causes a "hypertrophic myofibrillar cardiomyopathy". Journal of Muscle Research and Cell Motility. 10.1007/s10974-021-09601-1.
42. Ader, F., De Groote, P., Réant, P., Rooryck-Thambo, C., Dupin-Deguine, D., Rambaud, C., Khraiche, D., Perret, C., Pruny, J.F., Mathieu-Dramard, M., et al. (2019). FLNC pathogenic

variants in patients with cardiomyopathies: Prevalence and genotype-phenotype correlations. Clinical Genetics *96*, 317-329. 10.1111/cge.13594.

43. Yang, J., and Zhang, Y. (2015). I-TASSER server: new development for protein structure and function predictions. Nucleic Acids Research *43*, W174-W181. 10.1093/nar/gkv342.
